# The arthropod associates of 155 North American cynipid oak galls

**DOI:** 10.1101/2022.04.26.489445

**Authors:** Anna K.G. Ward, Robert W. Busbee, Rachel A. Chen, Charles K. Davis, Amanda L. Driscoe, Scott P. Egan, Bailey A.R. Goldberg, Glen Ray Hood, Dylan G. Jones, Adam J. Kranz, Shannon A. Meadely Dunphy, Alyson K. Milks, James R. Ott, Kirsten M. Prior, Sofia I. Sheikh, Shih-An Shzu, Kelly L. Weinersmith, Linyi Zhang, Y. Miles Zhang, Andrew A. Forbes

## Abstract

The identities of most arthropod associates of cynipid-induced oak galls in the western Palearctic are generally known. However, a comprehensive accounting of associates has been performed for only a small number of the galls induced by the estimated 700 species of cynipid gall wasp in the Nearctic. This gap in knowledge stymies many potential studies of diversity, coevolution, and community ecology, for which oak gall systems are otherwise ideal models. We report rearing records of insects and other arthropods from more than 527,306 individual galls representing 201 different oak gall types collected from 32 oak tree species in North America. Of the 201 gall types collected, 155 produced one or more animals. A total of 151,075 animals were found in association with these 155 gall types, and of these 61,044 (40.4%) were gall wasps while 90,031 (59.6%) were other arthropods. We identified all animals to superfamily, family, or, where possible, to genus. We provide raw numbers and summaries of collections, alongside notes on natural history, ecology, and previously published associations for each taxon. For eight common gall-associated genera (*Synergus*, *Ceroptres*, *Euceroptres*, *Ormyrus*, *Torymus*, *Eurytoma*, *Sycophila*, and *Euderus*), we also connect rearing records to gall wasp phylogeny, geography, and ecology - including host tree and gall location (host organ), and their co-occurrence with other insect genera. Though the diversity of gall wasps and the large size of these communities is such that many Nearctic oak gall-associated insects still remain undescribed, this large collection and identification effort should facilitate the testing of new and varied ecological and evolutionary hypotheses in Nearctic oak galls.

## Introduction

The biology of insect-induced plant galls has fascinated scientists for centuries and continues to provide a productive crucible for developmental, evolutionary, ecological, natural history, and applied studies (Darwin 1875, Fagan 1918, Weis and Abrahamson 1987, Price et al. 1987, Fernandes et al. 2014, Tooker and Helms 2014). Second only to the cecidomyiid gall midges (Diptera) in species richness (Ronquist & Liljeblad 2001), the oak galls wasps (Hymenoptera: Cynipidae: Cynipini) have been particularly well studied in many respects (reviewed by Stone et al. 2002) for three simple reasons: many of their galls are relatively large and conspicuous; galls are found on common and widespread tree species (mostly oaks); and inspection of each gall provides a wealth of information on both the fate of these sedentary insect herbivores and the interactions within and among associated gall occupants. However, though oak gall wasps have long been the subject of interest, we still know surprisingly little about some fundamental aspects of their biology. Some examples: the mechanism by which galls are induced has until recently been largely elusive (though see Hearn et al. 2019, Martinson et al. 2021); their taxonomy has been confounding and has frequently required revision at both large (e.g., Melika et al. 2021ab) and small scales (Zhang et al. 2021); many species remain undescribed (Sonte et al. 2002); and the life cycles of many species remain elusive, making it difficult to explore basic aspects of their ecology and life history. These difficulties are especially prominent in the Nearctic region, despite this being the home to the majority of global oak gall wasp and oak species diversity (Penzes et al. 2018, Melika et al. 2021b) and a rich historical legacy of collecting beginning with the pioneering works of Ashmead (1887, 1903), Weld (1921), and Kinsey (1923, 1930).

An additional and important aspect of Nearctic oak gall biology that is heretofore unknown and is explored herein is an understanding of both the composition and drivers of the diversity of the communities of insects and other arthropods that associate with gall wasps and their galls. For example, in the western Palearctic, the insect associates of many oak galls are well-studied, including, in some cases, their interactions and ecological roles (Askew 1961, Askew et al. 2013). The ∼150 western Palearctic gall wasps are associated with ∼33 endemic oak species (Denk and Grimm 2010) and the galls on those trees are host to at least 100 described wasp species and >30 inquiline cynipid wasp species (Askew et al. 2013). Some of these gall associates have been the subjects of ambitious and fruitful ecological and evolutionary studies (Stone and Schönrogge 2003, Bailey et al. 2009, Nicholls et al. 2018). However, in the Nearctic region, descriptions of arthropod communities associated with oak galls have lagged behind those in the western Palearctic in spite of (or more likely because of) the far more species-rich assemblages of both gall wasps and oak species in the Nearctic. The Nearctic is home to more than 150 species of oak (Hipp et al. 2018, Manos and Hipp 2021), which together are host to more than 700 of an estimated 1000 global oak gall wasp species (Melika et al. 2021b). In sum, the Nearctic gall wasps and their oak tree hosts promise a far more complex landscape for ecological, population genetic, phylogenetic, evolutionary, and functional studies across multiple trophic levels (and indeed this promise has already been kept: e.g., Prior and Hellmann 2013, Hood et al. 2019, Zhang et al. 2019), but such studies would greatly benefit from a more complete understanding of their associated insect communities.

Previous characterizations of the communities associated with Nearctic oak galls have generally taken a one-gall-at-a-time approach, involving collections of a single gall type and identification of emergent insects to the lowest achievable taxonomic level. In almost all cases, these studies have revealed diverse arthropod communities, with species numbers associated with individual gall types ranging from 15 to 25 (Joseph et al. 2011, Bird et al. 2013, Prior and Hellmann 2013, Forbes et al. 2016, Weinersmith et al. 2020). This approach provides a detailed inventory of the particular species associated with any specific type of gall and forms the basis for addressing intraspecific, and interspecific interactions within and among gall associate species and within and among trophic levels. However, for those interested in a regional level understanding of the ecological and evolutionary dynamics of oak gall wasps and their insect associates throughout the Nearctic, an overall picture synthesized from studies of individual gall types will take decades to produce. For that reason, we here take a many-galls-together approach, reporting a large number of previously undocumented associations.

Three developments now position researchers working in North America to explore fundamental dimensions of gall-associated insect diversification and together raise the possibility that some gall-associated insects may have diversified either via co-speciation with gall wasps or by frequently shifting between different host galls. One, the publication of a robust phylogeny of North American Cynipid oak gallers (Ward et al. 2022) now offers opportunities to contextualize insect gall associations across an evolutionary dimension. Two, several of the same insect natural enemy genera are known to be commonly reared from many different oak galls, and particular clades within each genus are known to specialize on oak galls (Gillette 1896, Balduf 1932, Grissell 1976, Hanson 1992, Lobato-Vila et al. 2019). Three, studies employing DNA barcoding and sequencing of Ultraconserved Elements (UCE) paired with ecological data show more diversity, and more specialization on gall hosts, than had previously been suggested for some taxa (Ward et al. 2020, Sheikh et al. 2022, Zhang et al. 2022). The data compiled and presented herein constitute a necessary step toward addressing coevolutionary (and other) hypotheses related to interactions between oak gall wasps and their insect associates.

Here, we report insect and other arthropod associates of 201 distinct types of oak gall collected from 32 oak species across continental North America. For 155 of these gall types, we report at least one, and as many as 21 associated insects or other arthropod species. The taxa reared from these collections span three classes, nine insect orders, and more than 29 insect families, variously acting as parasites, inquilines, hyperparasites, kleptoparasites, and successional associates in these gall systems. We provide a full summary table of all collection and rearing data to aid other researchers in planning their own collections. For eight of the most common genera of oak gall associates, we also summarize rearing records as they relate to gall wasp phylogeny and geography, as well as ecological data - including the host tree and gall location (host organ), and co-occurrence with other genera. These summaries provide insights into the myriad ways that different gall associates interact with galls and promise to be foundational in future evaluation of evolutionary and ecological hypotheses in these systems.

## Methods

Collections from three different “teams” are summarized in this paper. One set (authors AAF, AKGW, and SIS) was collected from 2015-2019 across much of the continental United States, but with an emphasis on the Midwest and Northeast. For these collections (identified in Suppl. Table 1 with three-digit numerical codes), galls of each type from each individual collection event were pooled and put into a plastic deli container with a mesh cover, and then held in a climate controlled incubator that mimicked seasonal changes in temperature, humidity, and light:dark cycles typical of the upper Midwest. Cups were checked once daily for emergent organisms, which were removed and put into individually labeled microcentrifuge tubes half-filled with 95% ethanol.

A second team (authors KMP, DGJ, RAC, AKM, and SAM) focused on galls on *Quercus garryana* Douglas ex. Hook in northern California, Oregon, Washington, and on Vancouver Island, British Columbia. These collections (alphabetical lab codes, Supp. Figure 1) were systematically collected at 10 oak patches (WA, BC) in 2017 and 18 oak patches in 2019 (CA, OR, WA, BC). Four (2017) and three (2019) collections were made between mid-May and early August at each site on a rotating basis, starting with southern sites. For each collection period, the team surveyed 10 trees per site (400 trees in 2017, 580 in 2019) and searched for and collected galls on ∼1 m of 10 branches. Galls were housed in plastic containers with mesh, separated by species, site and collection period. Containers were kept in environmental chambers set to Pacific Northwest summer conditions (25°C, 14:10) in a USDA-rearing room at Binghamton University. Emergent animals were collected once weekly for almost one year. Non-hymenopteran insects were not formally tallied for these collections and removed from collections when possible. However, lepidopteran and coleopteran adults and larvae were observed to emerge from large and fleshy galls induced by *Neuroterus washingtonensis* Beutenmüller and *Andricus quercuscalifornicus*.

**Figure 1.**
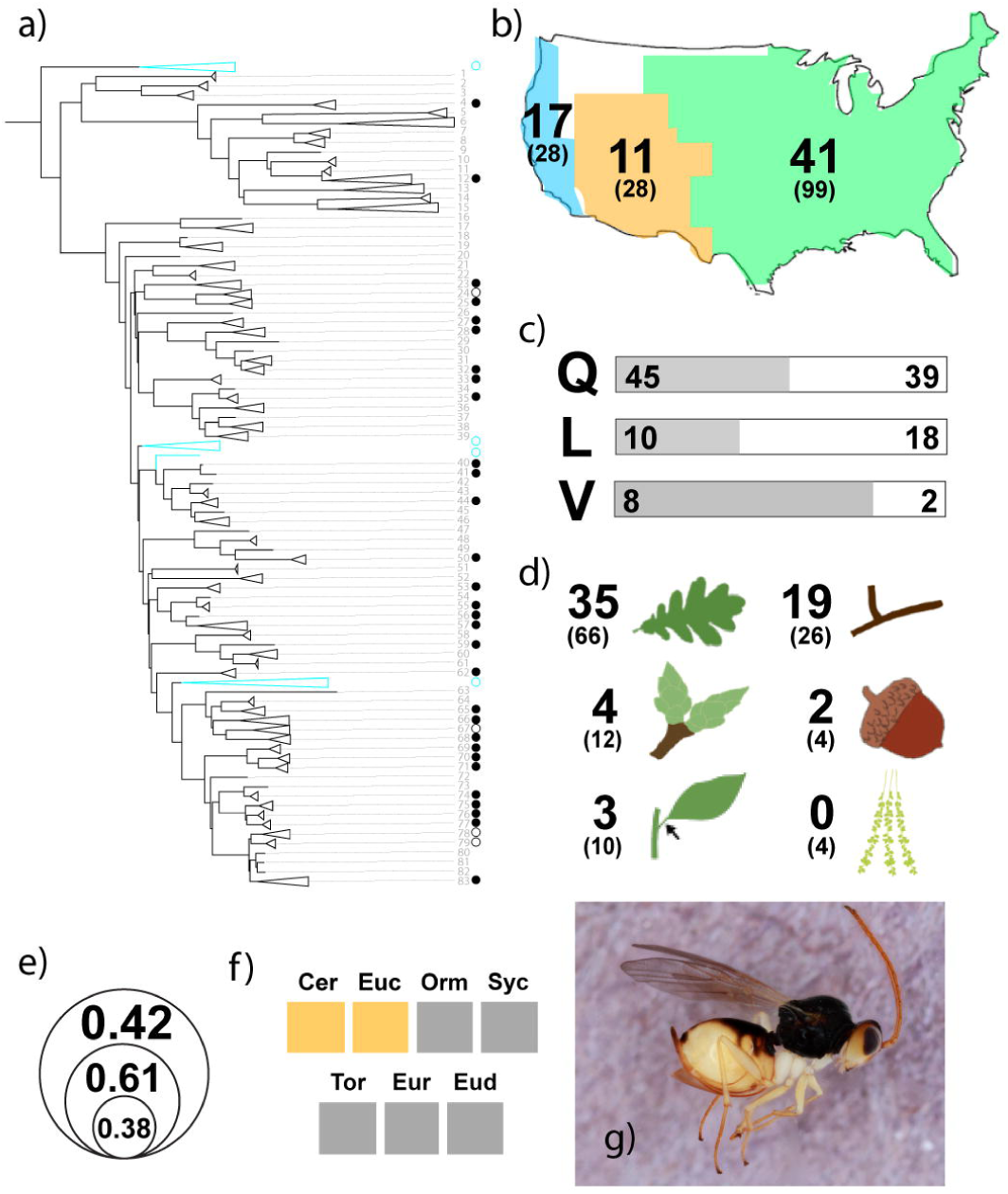
Summary of data for *Synergus* inquilines reared from Nearctic galls. a) Associations of *Synergus* mapped to the Ward et al. (2022) Nearctic oak gall wasp phylogeny (Supplementary Figure 1). Numbers at tips of branches refer to those in Supplementary Figure 1. Closed circles at branch tips indicate *Synergus* was reared from galls of that gall wasp species in this study. Open circles indicate other previously known associations either not studied by us or not recovered in our collections. Blue-colored branches within the phylogeny indicate Palearctic gall wasps. b) Total number of gall types from which *Synergus* were reared in the three bioregions identified by Hipp et al. (2018) as constituting different assemblages of North American oaks: Californian (blue), Mexican and Central American (orange), and Eastern North American (green) floristic provinces. Numbers in parentheses indicate the total number of gall types collected in each region, excluding gall types from which no insects emerged. For figures c-f, gall types from which fewer than five individual insects were reared were excluded, whether or not a *Synergus* was reared. c) Associations of *Synergus* with trees in sections *Quercus* (Q), *Lobatae* (L), and *Virentes* (V). Gray bars and numbers indicate gall types with which a *Synergus* was associated. White bars and numbers indicate the number of gall types from which a *Synergus* was not reared. d) Association of *Synergus* with gall types on different oak tissues (“Bud” includes galls that may be found on both buds or stems. “Petiole” includes galls that may be found on both petioles and stems or petioles and leaves); e) proportion of gall types of three size categories (“small” <0.5 mm; “medium”, “large” > 20 mm) from which *Synergus* were reared. f) Results of probabilistic co-occurrence analysis (Veech 2013) for *Synergus* against seven other common associates (Cer = *Ceroptres*, Euc = *Euceroptres*, Orm = *Ormyrus*, Syc = *Sycophila*, Tor = *Torymus*, Eur = *Eurytoma*, Eud = *Euderus*). Yellow = significantly less likely to co-occur; blue = significantly more likely to co-occur; gray = no difference from probabilistic expectations. g) *Synergus* lateral habitus.

The third group (authors JRO, SPE, RWB, ALD, GH, KLW, CKD, SS) was focused on collections exclusively from oaks in section *Virentes* in the south and southeastern United States. These galls (no lab code) were collected by hand in 2015-2019 from three species of live oaks: (1) *Quercus virginiana* Mill. distributed along the southeastern U.S. Gulf and Atlantic coasts from North Carolina south to Florida and west to Texas; (2) *Quercus geminata* Small, restricted to Florida, Mississippi, Georgia, and coastal Alabama; (3.) *Quercus fusiformis* Small, restricted to central and north-central Texas, as well as isolated populations in southwestern Oklahoma. In most cases, collections of multiple gall inducing species were made from multiple trees at a single site. These were separated by gall type and then housed in mason jars capped with inverted funnels connected to a *Drosophila* vial sealed with a cotton swab cap and housed at Texas State University (San Marcos, TX) and Rice University (Houston, TX) in the shade outdoors to match the light-dark, temperature and humidity cycles of the region. Containers were misted with deionized water once a week to maintain relative humidity levels that induce emergence. Collections were checked once every two days and emergent individuals were preserved by date, gall type, and collection locality. Some stem galls were placed in plastic bags and transported to either Rice University (Houston, TX) or Charlottesville, VA. Leaves, nontarget galls, and any invertebrates found on the exterior of the stems were removed, and stems were placed in clear plastic cups. The cups were covered with a coffee filter secured in place with a rubber band. Cups were then placed outside and treated as above.

Many cynipid wasps galling oaks undergo heterogony or cyclical parthenogenesis (Pujade-Villar et al. 2001), whereby asexual (agamic) and sexual (gamic) generations alternate yearly to complete a bivoltine life cycle. In most cases, the galls induced by the respective generations, which differ in morphology, are induced on a different organ within the same host plant. Thus, the two generations typically induce galls on different host organs at different times of the year. Where we made collections from both the gamic and agamic galls induced by the same gall wasp species, we counted these as different gall hosts, reflecting that the morphology, oak organ galled, and the timing of gall formation are almost always different and thus likely to harbor communities of gall associates that differ (e.g. Forbes et al. 2016). For this reason, when referring to the unit from which insects emerged, we use “gall types” rather than “gall former species.” Galls (and by proxy, the species that induce the gall type) were identified based on gall morphology (Weld and von Dalla Torre 1952, Weld 1957, 1959, 1960, Melika & Abrahamson 2002, Russo 2021, “gallformers.org” 2022). We refer to some unnamed galls by their descriptions on gallformers.org as of January 2022, in Russo 2021, or else by our own descriptor when no similar gall was found in the literature or on the gallformers.org website.

We keyed all emergent arthropods to superfamily, family, or, where possible, to genus (McAlpine et al. 1981, Goulet and Huber 1993, Gibson et al. 1997, Arnett et al. 2000, 2002). Though species-level keys based on morphology exist for some Nearctic gall-associated genera, recent molecular analyses of three genera – *Synergus* Hartig, *Ormyrus* Westwood, and *Sycophila* Walker – have revealed much cryptic diversity in each genus (Ward et al. 2020, Sheikh et al. 2022, Zhang et al. 2022), echoing similar work in the Palearctic (Ács et al. 2002, Kaartinen et al. 2010). Thus, to avoid incorrectly ascribing host associations of several putative specialist species to a single “lumped” species on the basis of morphology alone, we intentionally did not key many collections to species. Only in the cases of the relatively species-poor genera *Euderus* Halliday and *Euceroptres* Ashmead did we key individuals to species, though even here we suggest caution in interpreting species-level host ranges. Due to the large size of these collections and our intention to use many in various future projects, we have not yet made a full set of voucher specimens available for study. However, vouchers for many of our gall wasps, and for wasps in the genera *Synergus*, *Ormyrus,* and *Sycophila* are already available (Ward et al. 2020, Sheikh et al. 2022, Ward et al. 2022, Zhang et al. 2022) and we intend to make vouchers of other genera available as more publications emerge from this dataset. Researchers interested in this material sooner than vouchers are made available may contact authors AAF, SPE, or KMP.

For eight of the most commonly collected genera (*Synergus*, *Ceroptres* Hartig, *Euceroptres*, *Ormyrus, Torymus* Dalman, *Sycophila*, *Eurytoma* Illiger, and *Euderus*), we had a sufficient collection to be able to ask some common questions about their associations. We used a common treatment to this end:

a) Some insect genera might be associated with a phylogenetically-limited subset of gall wasps, and, if so, such patterns might inform hypotheses about the ecology and evolution of those gall associates. Thus, we created a slightly modified version (Supplemental Figure 1) of a recently published North American gall wasp phylogeny (Ward et al. 2022), and then mapped collection data for each member of the genus onto that tree. We also mapped historical records of collections onto the tree when those records were available (Gillette 1896; Krombein et al. 1979; Bugbee 1967; Yoshimoto 1971; Hansen 1992; Noyes 2022).
b) Some insect genera may be geographically restricted, reflecting their particular biogeographic histories. Thus, we divided our collection data into the three floristic provinces for oaks used by Hipp et al. (2018): Californian (CA), Mexican & Central American (MCA), and Eastern North American (ENM) to visualize geographic patterns of host associations for each genus and province. For these visualizations we excluded gall types from which no insects emerged. Total gall numbers in map figures are inflated by one because one gall type (*Andricus quercuslanigera* (Ashmead)) was found in both ENM and MCA and was counted separately in both.
c) Each oak gall wasp species specializes on a taxonomically-restricted set of oak trees (Ward et al. 2022). To determine whether other insects might also be restricted to certain trees, we arranged data based on each genus’ association with oaks in three sections: *Quercus* (white oaks), *Lobatae* (red oaks), and *Virentes* (live oaks). Because collection and rearing methods may have impeded insect development in some galls, we only included data from galls from which we had reared at least five arthropods of any type.
d) Just as insect genera might specialize on one or more oak tree sections, they could favor galls in specific locations on the tree. Thus, we organized data by the type of oak organ with which host galls were associated. Again, we only included data from galls from which we had reared at least five insects of any type.
e) Some insect genera associated with galls might prefer, or require for development, galls in a particular size range. Thus, we organized data based on gall size (“small” < 5 mm; “medium” > 5 mm & < 20 mm, “large” > 20 mm), again using only data from galls from which we had reared at least five insects of any type.
f) Finally, some gall associates may require presence of other genera (i.e, if they are obligate hyperparasitoids or parasites of inquilines) or might converge with other genera in attacking galls with similar characteristics. Therefore, for the 47 gall types from which we reared more than 100 non-gall wasp insects, we conducted a probabilistic co-occurrence analysis (Veech 2013). We converted collections data into presence/absence data and tested for positive and negative co-occurrence among eight common associates using the R package *co-occur* (Griffith et al. 2016), filtering out any pairs from the dataset that would be expected to share >1 galler based on a probabilistic model.

Beyond these eight common genera, we report numbers and general patterns of host gall association for one superfamily (Ichneumonoidea) and three commonly-reared families (Eulophidae, Eupelmidae, and Pteromalidae), that we could not confidently separate to genus. We also report records of a large number of insects and other arthropods from diverse groups that, for shared or different reasons, were reared from a relatively smaller fraction of collected galls.

### Caveats and Omissions

The data we report below are complex and imperfect in several ways that bear careful consideration:

#### Negative data

While a confirmed rearing event documents an association of an insect with a particular type of gall, failure to rear an insect from a gall type cannot be interpreted as a lack of association in nature as there are many reasons why any given associate might be present but not reared. For instance, the timing of our removal of galls from trees may cause some galls to prematurely desiccate or otherwise reduce the quality of the resource they provide to insects inside, such that some insects that would have emerged in the wild do not emerge in the laboratory setting. Similarly, detachment of galls from the host plants and or rearing conditions in the lab may not provide appropriate plant mediated or abiotic environment mediated developmental triggers for some insects, such that they do not emerge as readily as they would in the field.

Similarly, spatial and temporal variation within and among our collections almost certainly affected the composition of the assemblage of emergent insects. Some galls may have been collected before the developmental stage at which a particular associate uses the gall or collected after that associated insect has emerged from the gall. Some associates also may not overlap completely with the geographic range of a gall. With the exception of collections of galls induced by the three species of *Belonocnema* Mayr on live oaks (Busbee 2018; Driscoe et al. 2019), none of our collections covered the entirety of any gall wasp’s geographic range in the U.S.A., nor were any galls collected systematically from gall induction until effective decomposition.

#### Omitted gall types

These collections inevitably favored more prominent galls, galls that are more common and or abundant, and galls that persist on plants for longer periods. All galls in this study were aboveground (i.e., we collected no galls from oak roots, though many are known; Weld 1965). Similarly, galls that are concealed under bark with no obvious external swelling were not collected in this study.

#### Lumping / splitting of gall wasp species

Most gall wasp species have been described based solely on their adult morphology. Molecular analyses have since identified genetic differentiation, often correlated with differences in host tree species, for several Nearctic gall wasp species (Cooke 2018, Drisccoe et al. 2019, Ward et al. 2022) such that many “gall wasp species” may actually constitute one or more different species. Extending this problem further: some of the “communities” compiled here could therefore represent conglomerates of several communities, each with different members.

#### Diversity / Species Richness measures

Though a large collection of insects sampled from many different communities and guilds seems appropriate for calculating various metrics of richness and diversity, we do not provide these as our reporting of gall associates herein spans several different scales of taxonomic organization (order, superfamily, family, genus, species). Thus, calculating these metrics is premature pending species designations and would be of little obvious utility. The haphazard nature of many of our collections, the variation in numbers of individuals collected for each gall type, as well as some of the same caveats listed above also preclude us from being able to estimate and compare diversity.

With those caveats, what follows is a report of a large collection of oak gall-associated arthropods, contextualized ecologically, geographically, and (at least in a preliminary way) evolutionarily. This report sets the stage for more incisive future studies.

## RESULTS AND DISCUSSION

Across 1,789 collection events, we accumulated more than 527,275 individual galls representing 201 different oak gall types collected from 32 oak tree species. Across all collections a total of 151,075 animals emerged (61,044 gall wasps, 90,031 other arthropods). Supplementary Table 1 summarizes collections and rearing records by gall type. Forty-seven gall types yielded no animals, though these were typically low-number gall collections (range 1–154 galls; median = 9). From the remaining 155 gall types, between 1 and 70,435 animals were reared (median = 33) from collections totaling from 1 to 254,171 galls (median = 47). At least five animals emerged from each of 122 gall types. Co-occurrence analysis results showed three significant negative associations and four significant positive associations among gall-associated insect genera. Other relationships did not differ significantly from probabilistic expectations. Outcomes of co-occurrence analyses are shown for each genus in Figures 1–8 and in Supplementary Table 2. Below we discuss each type of gall associate individually, starting with the gallers themselves, followed by the eight most commonly identified genera, then four other families or superfamilies not identified to the genus level, and finally by other less common associates.

### Gall wasps (Hymenoptera: Cynipoidea: Cynipidae: Cynipini)

Gall-inducing wasps (“gall wasps”) accounted for 61,044 (40.4%) of all emerging animals. No gall wasps emerged from 99 of the 201 gall types, including from some gall types that were collected in relatively high numbers (range: 1–2,790 galls; median: 17 galls). Fifty three of the 99 gall types that bore no gall wasps had other arthropods emerge. Six gall types produced gall wasps and no other insects; these were all relatively small collections, ranging from just 1 to 35 total galls collected.

#### *Synergus* Hartig 1840 (Hymenoptera: Cynipoidea: Cynipidae: Synergini)

7,227 individuals (mean = 104.7, range 1 – 3,228) reared from 69 gall types (Supplementary Table 1).

#### Summary of Natural History

*Synergus* (Hymenoptera: Cynipidae: Synergini) are usually professed to be inquilines, but are perhaps more accurately described as gallers of galls (Askew 1961). *Synergus* induce additional growth in existing galls, including the formation of larval chambers, and their developing young feed on the tissue of the gall (Evans 1965). Though gall inducing *Synergus* have been documented in Japan (Abe et al. 2011, Ide et al. 2018), gall induction in *Synergus* is a derived habit (Ide et al. 2018) and not known from the Nearctic. In some galls, the presence of *Synergus* is fatal to the developing gall inducer, but in other cases food may be sufficient such that both may emerge (Pénzes et al. 2012). In some galls, *Synergus* develop and emerge as adults within a matter of weeks, while others can take one or even two years to emerge (Evans 1965, Busbee 2018, Ward et al. 2020).

Multiple species of *Synergus* can be associated with the same gall type (Askew 1961; Penzes et al. 2012; Bird et al. 2013; Forbes et al. 2016, Weinersmith et al. 2020), while other galls have no known *Synergus* associates despite large collection efforts (e.g., Joseph et al. 2011). There has been some previous suggestion that two other genera of cynipid inquilines (*Ceroptres* and *Euceroptres*) of Nearctic gall wasps may not co-occur with *Synergus* (Brookfield 1972), but curated rearing records (e.g., Krombein et al. 1979) and our own data presented here show that this is not universally true.

#### Relationship to galler phylogeny

*Synergus* wasps were reared from gall types across most of the Nearctic gall wasp phylogeny (Ward et al. 2022), with some exceptions. In only two cases were *Synergus* reared from gall types produced by gallers in the large clade that includes genera *Melikaiella* Pujade-Villar, *Loxaulus* Mayr, and most of the *Neuroterus* Hartig (Supplementary Figure 1; gallers #2-15 in Figure 1a). Both of these *Synergus* / *Neuroterus* associations were from Pacific coast galls. The reduced apparent association of *Synergus* with gall inducers in this clade might reflect that the *Synergus* association with oak gall wasps originated in the clade represented by the lower two-thirds of the tree.

However this hypothesized relationship is belied by records of *Synergus* being associated with the topmost clade of Palearctic gall wasps in Figure 1a (though these could represent secondary colonizations). Ultimately, assessment of coevolutionary relationships of *Synergus* with oak gall wasps requires a phylogeny of the Holarctic *Synergus*.

#### Biogeography and oak tree section

*Synergus* were reared from galls in all three North American oak floristic regions and from galls on trees across all three sampled oak sections (Figure 1b,c). Of the three sections, *Synergus* were least often reared from galls on section *Lobatae*.

#### Tree organ and gall size

*Synergus* were reared from galls developing on leaves, stems, buds, acorns, and petioles (Figure 1d). Among organs from which we sampled galls, only flower galls did not have apparent *Synergus* associates. *Synergus* in our collections were most commonly reared from medium sized galls (61%) and least commonly reared from small galls (38% of galls smaller than 5 mm) (Figure 1e). Though these differences are not large, they comport with observations of Palearctic *Andricus* Hartig galls which suggest that small bud and catkin galls were less likely to host *Synergus* (Stone et al. 1995). Alternatively, reduced association of *Synergus* with small bud and catkin galls could be related to their earlier temporal occurrence: 25 of the 27 putative *Synergus* species in Ward et al. (2020) were reared from galls developing in June or later. However, we again raise the caveat that small galls may desiccate in the lab causing associated insects to die before emergence and leading to apparent non-associations.

#### Co-occurrence with other natural enemies

When *Synergus* were present, two other putative inquiline genera were significantly less likely to be present: *Euceroptres* (P = 0.005) and *Ceroptres* (P = 0.0026). Thus while our data disagree with the suggestion that *Synergus* and *Ceroptres* / *Euceroptres* entirely displace one another (Brookfield 1972), they do appear to co-occur less often than expected. This could be due to competitive exclusion, but also or instead be an indirect result of differential adaptation to dimensions of gall hosts. Notably, different *Synergus* species do not apparently competitively exclude one another, with as many as five species having been reared from the same host gall (Penzes et al. 2012). To the extent that more closely related species are expected to compete more closely when sharing the same habitat (Miller 1967, Denno et al. 1995), differential adaptation to some dimensions of the gall environment seems the more attractive hypothesis.

#### Additional notes

Our record of four *Synergus* wasps reared from galls of *Andricus quercuscalifornicus* appear to be the first ever, despite much attention having been paid to this particularly large and common Pacific coast gall and its natural enemies (Joseph et al. 2011). Other efforts to collect and rear insects from large numbers of potential hosts often turn up uncommon associations (Yee 2008, Yee and Goughnour 2008). Given that host shifts have often been implicated in the origins of parasitic insect diversity (Diehl and Bush 1984, Drés and Mallet 2002, Forbes et al. 2017), evidence of insects occasionally being reared from unexpected hosts suggest that variation in host recognition syndromes may result in insects often “testing” new potential host plants.

#### *Ceroptres* Hartig 1840 (Hymenoptera: Cynipoidea: Cynipidae: Ceroptresini)

4,335 individuals (mean = 88.5, range 1 – 1,683) reared from 49 gall types (Supplementary Table 1).

#### Summary of Natural History

The genus *Ceroptres* are oak-associated putative inquilines (or kleptoparasitic gall-modifiers; Ronquist 1994), but the biology of members of this genus is not as well studied as many *Synergus*, therefore their role requires further investigation. Originally in the same tribe as *Synergus*, they have been moved to their own tribe, the Ceroptresini, reflecting their likely independent evolution of inquilinism (Nylander 2004; Ács et al. 2010). *Ceroptres* have a Holarctic distribution with most named species from the Nearctic (Penzes 2012; Lobato-Vila and Pujade-Villar 2019), though it is probable that more species remain to be described from the Eastern Palearctic (Wang et al. 2012). Note that our “*Ceroptres*” does not distinguish between *Ceroptres sensu stricto* and the recently described and currently monotypic genus *Buffingtonella* (Lobato-Vila and Pujade-Villar 2019)

#### Relationship to galler phylogeny

*Ceroptres* were associated with gall inducers across the Nearctic gall wasp phylogeny (Ward et al. 2022), and are also known from Palearctic galls, including the most basal on the tree (Figure 2a). *Ceroptres* were not reared from some large gall wasp clades, including 1) the *Amphibolips* Reinhard galls, 2) a clade of *Andricus* and *Callirhytis* Förster bud and leaf galls, and 3) a clade of primarily Pacific coast galls (though no *Ceroptres* were reared from any Pacific coast galls).

**Figure 2.**
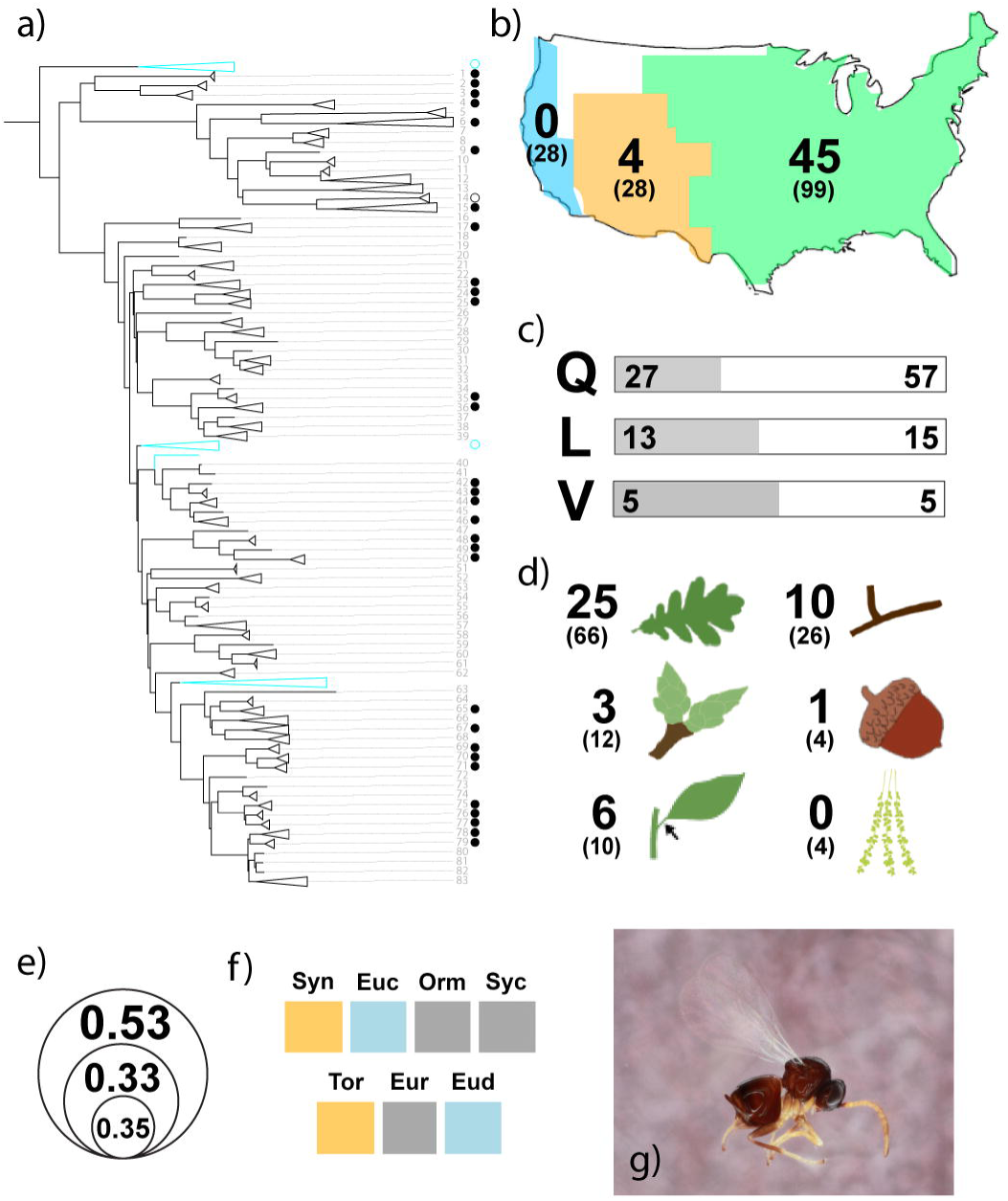
Summary of data for *Ceroptres* inquilines collected from Nearctic galls. For full explanation of figure details, refer to Figure 1 legend. a) Associations of *Ceroptres* with the Nearctic oak gall wasp phylogeny (Supplementary Figure 1). b) Gall types from which *Ceroptres* were reared in the Californian (blue), Mexican and Central American (orange), and Eastern North American (green) floristic provinces. c) Associations of *Ceroptres* with trees in sections Quercus (Q), Lobatae (L), and Virentes (V). d) Association of *Ceroptres* with gall types on different oak tissues. e) Proportion of gall types of three size categories (“small” <0.5 mm; “medium”, “large” > 20 mm) from which *Ceroptres* were reared. f) Results of probabilistic co-occurrence analysis for *Ceroptres* against seven other common associates. Yellow = significantly less likely to co-occur; blue = significantly more likely to co-occur; gray = no difference from probabilistic expectations. g) *Ceroptres* lateral habitus.

*Ceroptres* were also never reared from any of the more than 13,000 *Belonocnema* Mayr galls collected from across the southeastern U.S.A.

#### Biogeography and oak tree section

*Ceroptres* were reared much more commonly from galls collected in the Eastern half of the United States than in the Southwest or the Pacific coast (Figure 2b). We reared no *Ceroptres* from any Pacific coast gall types, though at least two species are known from California (McCracken and Egbert 1922). Galls from all three focal oak sections produced *Ceroptres*, with the smallest proportion from Section Quercus (Figure 2c). *Ceroptres* are also known from oaks in section *Cerris* in the Palearctic and section Protobalanus in California (Lobato-Vila and Pujade-Villar 2019).

#### Tree organ and gall size

*Ceroptres* emerged from galls on leaves, stems, buds, acorns and petioles, but not from flower galls (Figure 2d). A greater fraction of large (>20mm) gall types produced *Ceroptres* than small or medium sized galls (Figure 2e).

#### Co-occurrence with other natural enemies

*Ceroptres* had more significant correlations (four) with other gall associates than any other genus (Figure 2f). Two correlations were positive: *Ceroptres* co-occurred more often than expected with another inquiline genus, *Euceroptres* (see below), and with the parasitoid genus *Euderus*. *Euderus* have been suggested to specialize on galls without external spines or hairs (Ward et al. 2019), so it is possible that some species of *Ceroptres* have similarly restricted host ranges. Significantly less likely to co-occur with *Ceroptres* were *Synergus* (P = 0.0026) and *Torymus* (P = 0.0003) parasitoids (see discussion in those sections)

#### *Euceroptres* Ashmead 1896 (Hymenoptera: Cynipoidea: Figitidae: Euceroptrinae)

803 individuals (mean = 133.8, range 5 – 727) reared from six gall types (Supplementary Table 1).

#### Summary of Natural History

The biology of *Euceroptres* is not well studied and it is not known whether members of this genus function as inquilines, parasitoids, or hyperparasitoids. Only four species of *Euceroptres* are previously described from galls of five species of gall wasps (Buffington and Liljeblad 2008). All known hosts are oak galls. The apparently small number of species and hosts for *Euceroptres*, coupled with the possibility that each species may have a limited range of hosts has led to the speculation that these are the surviving members of a previously more species-rich genus (Buffington and Liljeblad 2008).

#### Relationship to galler phylogeny

Unlike the two genera of cynipoid associates of oak galls treated above, there are no published records of *Euceroptres* from the Palearctic (Buffington and Liljeblad 2008), though one unpublished record from Serbia is mentioned in Buffington et al. (2020). Most Nearctic gallers in our collections were not found to have associations with *Euceroptres*, though the nine that mapped onto the galler phylogeny were widely scattered across the tree (Figure 3a).

**Figure 3.**
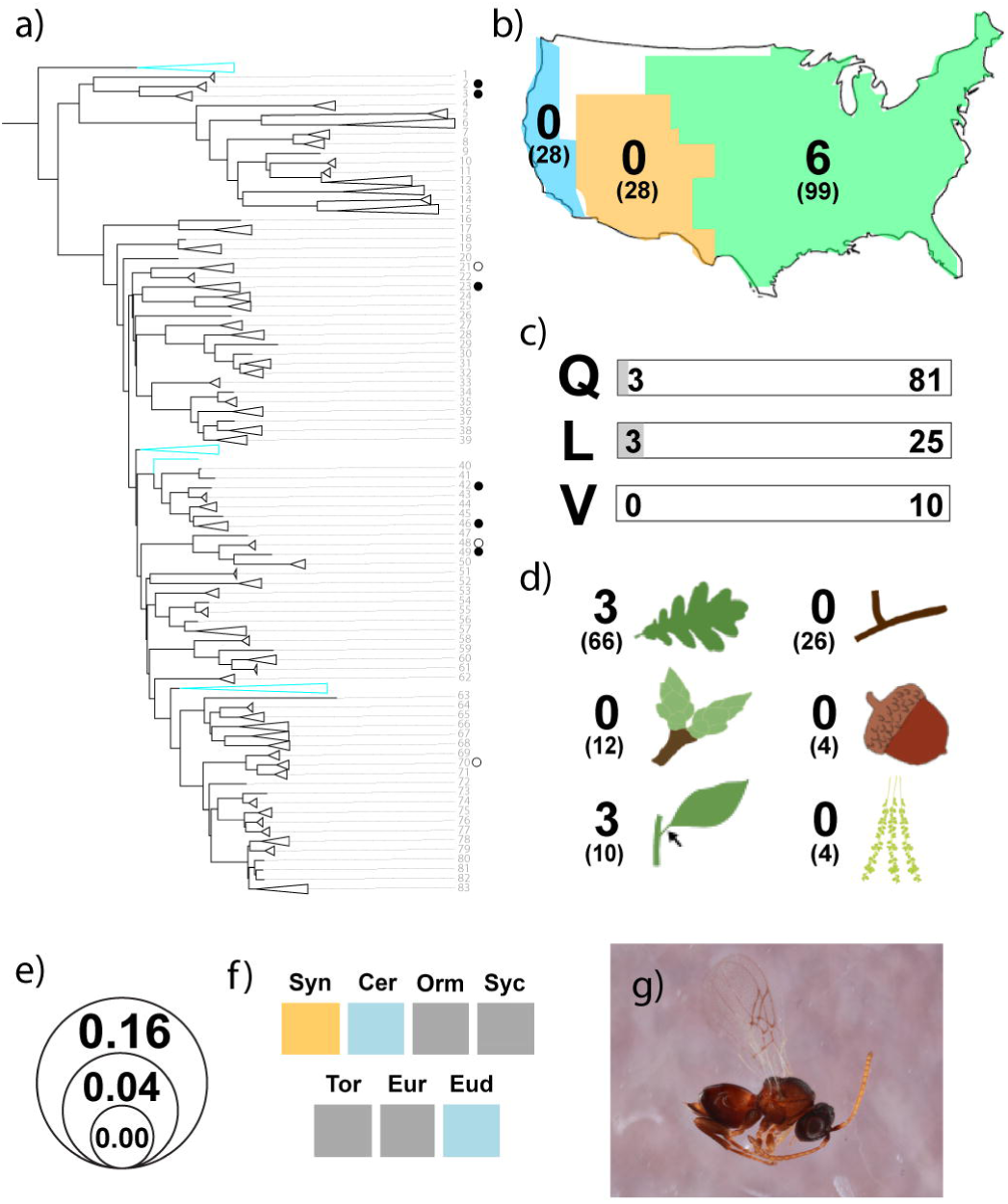
Summary of data for *Euceroptres* inquilines collected from Nearctic galls. For full explanation of figure details, refer to Figure 1 legend. a) Associations of *Euceroptres* with the Nearctic oak gall wasp phylogeny (Supplementary Figure 1). b) Gall types from which *Euceroptres* were reared in the Californian (blue), Mexican and Central American (orange), and Eastern North American (green) floristic provinces. c) Associations of *Euceroptres* with trees in sections Quercus (Q), Lobatae (L), and Virentes (V). d) Association of *Euceroptres* with gall types on different oak tissues. e) proportion of gall types of three size categories (“small” <0.5 mm; “medium”, “large” > 20 mm) from which *Euceroptres* were reared. f) Results of probabilistic co-occurrence analysis for *Euceroptres* against seven other common associates. Yellow = significantly less likely to co-occur; blue = significantly more likely to co-occur; gray = no difference from probabilistic expectations. g) *Euceroptres* lateral habitus.

#### Biogeography and oak tree section

All six species reared in this effort were from galls collected in the Eastern North American floristic region (Figure 3b). However, two species (*Euceroptres maritimus* Weld on *Callirhytis quercussuttoni* (Bassett) and *Euceroptres montanus* Weld on *Disholandricus truckeensis* (Ashmead)) were previously reared from California and Oregon. Three of the six gall hosts in our collections were collected from trees in oak section *Quercus*, while the remaining three were from trees in section *Lobatae* (Figure 3c). Previous collections were from galls found on oaks in sections *Quercus* and *Lobatae* as well as *Protobalanus* (Buffington and Liljeblad 2008; Manos and Hipp 2021). If *Euceroptres* are host specific as has been previously suggested (Buffington and Liljeblad 2008), our failure to rear them from western collections may well be a function of our collections not having included specific hosts.

#### Tree organ and gall size

All *Euceroptres* reared in this study were from galls on leaves or on petioles (Figure 3d). The host gall farthest from the leaf proper was *Callirhytis scitula* Bassett, a woody gall that occurs at the intersection between the petiole and stem. The two *Euceroptres* species known from the Pacific coast, however, are both associated with stem galls (Buffington and Liljeblad 2008). All six hosts among our collections were classified as either medium or large, with no rearings from any galls smaller than 5 mm (Figure 3e).

#### Co-occurrence with other natural enemies

*Euceroptres* were significantly more likely to co-occur with *Ceroptres* (P = 0.008) and with the parasitoid genus *Euderus* (P = 0.015) (Figure 3f). All six gall types from which we reared *Euceroptres* also had *Ceroptres* emerge and only one gall type from which we reared *Euceroptres* failed to yield *Euderus* (*Andricus foliaformis* Gillette). In contrast, *Euceroptres* were significantly negatively correlated with *Synergus* inquilines (P = 0.005) and in fact the two genera were never reared from the same gall types.

#### Additional Notes

The small number of rearing records for *Euceroptres*, both in the present study and historically, precludes definitive statements about this genus and its association with oak gall types. On the other hand, we now have initial information about which gall types do, and more importantly, apparently do not host *Euceroptres*. All six gall hosts identified here, and the few additional previously published host records (Buffington and Liljeblad 2008), are multi-chambered, integral galls (i.e., non-detachable) and larger than 0.5 cm, suggesting that one or more of these characters may be important with respect to host range for *Euceroptres*. However, many other galls with these same characters do not appear to be *Euceroptres* hosts, so while these characters may be necessary, it is not clear that they are sufficient for *Euceroptres* attack and development. It may be that *Euceroptres* requires the presence of another gall associate either because it is a parasite of wasps in one of these genera or because it is an inquiline or gall inducer that first requires, e.g., *Ceroptres* to make a gall within the gall (which would make *Euceroptres* a galler of galls-within-galls).

Though we generally did not key our collections to species, the small number of collections and the availability of the key to the four species produced by Buffington and Liljeblad (2008) allowed us to make an exception for *Euceroptres.* Not surprisingly, given the growing recognition of cryptic diversity and higher than previously-recognised host specificity among parasitic wasps (Forbes et al. 2009, Hood et al. 2015, Smith et al. 2011, Condon et al. 2014, Sheikh et al. 2022), none of the wasps we reared matched the descriptions of any of the four named Nearctic species. All samples had 10 flagellomeres and well developed micropores on their abdominal tergites, combinations not found in Buffington and Liljeblad (2008). None of the cynipid hosts producing the gall types from which we reared *Euceroptres* overlapped with previous rearing records, save for *Euceroptres whartoni* Buffington & Liljeblad in galls of *Andricus quercuspetiolicola* (Bassett). However, Buffington and Liljeblad (2008) list this host record as dubious.

Our collections to date may therefore include one or more undescribed species.

#### *Ormyrus* Westwood 1832 (Hymenoptera: Chalcidoidea: Ormyridae)

4,658 individuals (mean = 67.5, range 1 – 2,175) reared from 69 gall types (Supplementary Table 1).

#### Summary of Natural History

In comparison to other genera of parasitoids associated with oak galls, wasps in the genus *Ormyrus* have been described as generally more restricted to galls on oaks. For example, *Torymus* and *Eurytoma* both have oak gall associated species but attack many other hosts as well. All but two of the known Nearctic *Ormyrus* species attack galls on oaks – the exceptions attack Cynipid gall wasps on roses and Pteromalid gall inducers on blueberries (Hanson 1992). Though 16 species were described in the most recent revision of Nearctic *Ormyrus* (Hanson 1992), more recent genetic and ecological data suggest that these wasps are considerably more species rich and ecologically specialized than previously supposed (Sheikh et al. 2022).

*Ormyrus* larvae are generally known as ectoparasites, and in the Palearctic have been shown to directly parasitize the larvae of gall forming wasps, though it is also possible that they can attack inquilines and other parasitoids in the same galls (Redfern and Askew 1998) or that they are inquilines themselves.

Indeed, observations in the context of biocontrol suggest that *Ormyrus* can act as hyperparasitoids of *Torymus* wasps (Cooper and Rieske 2011). Moreover, experiments tracking insects emerging from individual *Belocnonema* leaf galls that are unilocular (or contain only one gall wasp larvae) show that *Ormyrus* wasps can emerge from the same individual gall as other adult gall associates, including *Synergus*, *Sycophila*, and *Brasema* Cameron (Hall 2001). Multiple individual *Ormyrus* were also observed emerging from the same individual gall (Hall 2001), which suggests that they were acting as inquilines or attacking insects other than the gall former.

#### Relationship to galler phylogeny

*Ormyrus* were reared in this study or previously reared by others from almost all gall types included in the Nearctic gall wasp phylogeny (Ward et al. 2022) phylogeny, including from all four Palearctic clades (Figure 4a). Members of the genus are near ubiquitous in associations with oak gall wasps.

**Figure 4.**
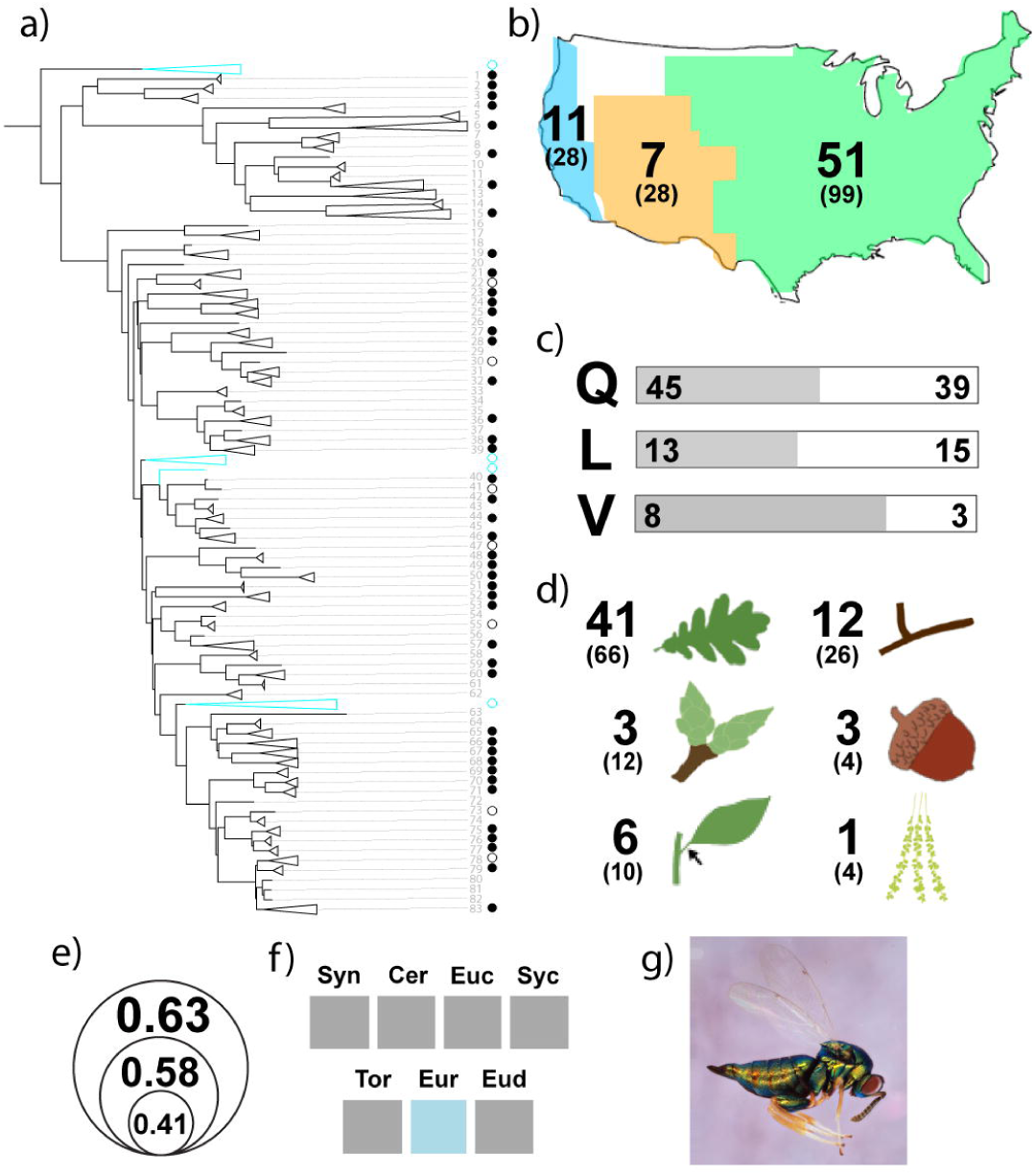
Summary of data for *Ormyrus* collected from Nearctic galls. For full explanation of figure details, refer to Figure 1 legend. a) Associations of *Ormyrus* with the Nearctic oak gall wasp phylogeny (Supplementary Figure 1). b) Gall types from which *Ormyrus* were reared in the Californian (blue), Mexican and Central American (orange), and Eastern North American (green) floristic provinces. c) Associations of *Ormyrus* with trees in sections Quercus (Q), Lobatae (L), and Virentes (V). d) Association of *Ormyrus* with gall types on different oak tissues. e) proportion of gall types of three size categories (“small” <0.5 mm; “medium”, “large” > 20 mm) from which *Ormyrus* were reared. f) Results of probabilistic co-occurrence analysis for *Ormyrus* against seven other common associates. Yellow = significantly less likely to co-occur; blue = significantly more likely to co-occur; gray = no difference from probabilistic expectations. g) *Ormyrus* lateral habitus.

#### Biogeography and oak tree section

*Ormyrus* were reared from galls collected in all three floristic regions of North America (Figure 4b) and from 45% or more of gall types monitored in each of the three oak tree sections surveyed (Figure 4c).

#### Tree organ and gall size

*Ormyrus* were associated with galls on all tree organs represented in our collections (Figure 4d) and from galls of all sizes, with a slight bias towards larger galls (Figure 4e). Previous rearing records also show that *Ormyrus* can use root galls produced by cynipids (Hanson 1992).

#### Co-occurrence with other natural enemies

We found only one significant correlation between *Ormyrus* and another insect genus: a positive correlation with *Eurytoma* (Figure 4f; P = 0.0033). This could represent a shared affinity for galls with similar characteristics or a currently unknown trophic association between *Ormyrus* and *Eurytoma* wasps (i.e., one is a parasite of the other).

#### Notes

As a genus, *Ormyrus* are ubiquitous in their phylogenetic, geographic, taxonomic, and ecological relationships with oak gall wasps. Though individual species are often apparently specialized on a small number of gall types (Sheikh et al. 2022), they may be able to use any of several insect species within a gall as their host, or act as inquilines, or some combination thereof. Understanding host breadths and life histories of individual *Ormyrus* species alongside their phylogeny will be critical to understanding how and under what circumstances oak-associated *Ormyrus* have diversified.

#### *Torymus* Dalman, 1820 (Hymenoptera: Chalcidoidea: Torymidae: Toryminae)

6,338 individuals (mean = 111.2, range 1 – 2,378) reared from 57 gall types (Supplementary Table 1).

#### Summary of Natural History

*Torymus* wasps are primarily Holarctic in their distribution and there are more than 300 named species, though not all of them attack oak gall wasps (Grissell 1995). Hosts are almost always insects in a concealed location, but from diverse orders. In the Nearctic, two species groups contain all of the oak gall-associated species. The *fullawayi* species group consists primarily of species that attack insects in cynipid oak galls (Grissell 1976). Species in the *tubicola* group have a more diverse range of hosts, but several specialize on oak galls or on galls on other plant hosts (Grissell 1976).

Most female *Torymus* have a long ovipositor (in some species, the ovipositor is more than the length of their body) that helps them attack hosts in concealed and otherwise protected locations. There is some evidence for apparent intraspecific variation in ovipositor length in some species, both between generations and within the same generation (Eady 1958, Askew 1965), which may result in a larger than expected host range for any given species. In the Palearctic, oak gall associated species range between one and 41 host records (Askew et al. 2013). The Nearctic species *Torymus tubicola* (Osten Sacken) is also described as a particularly generalist species, with more than 30 named oak gall hosts across the United States (Grissell 1976, Noyes 2022). However, as for *Ormyrus* and *Synergus* some of the apparently generalist *Torymus* species in both the Palearctic (Kaartinen et al. 2010) and Nearctic (Goldberg and Forbes, in prep; Driscoe et al., in prep) may harbor cryptic species.

#### Relationship to galler phylogeny

*Torymus* wasps have been reared from gallers across the Nearctic oak gall wasp phylogeny (Ward et al. 2022), including from all four Palearctic lineages (Figure 5a). Like *Ormyrus*, the genus houses essentially ubiquitous parasitoids in the oak gall system.

**Figure 5.**
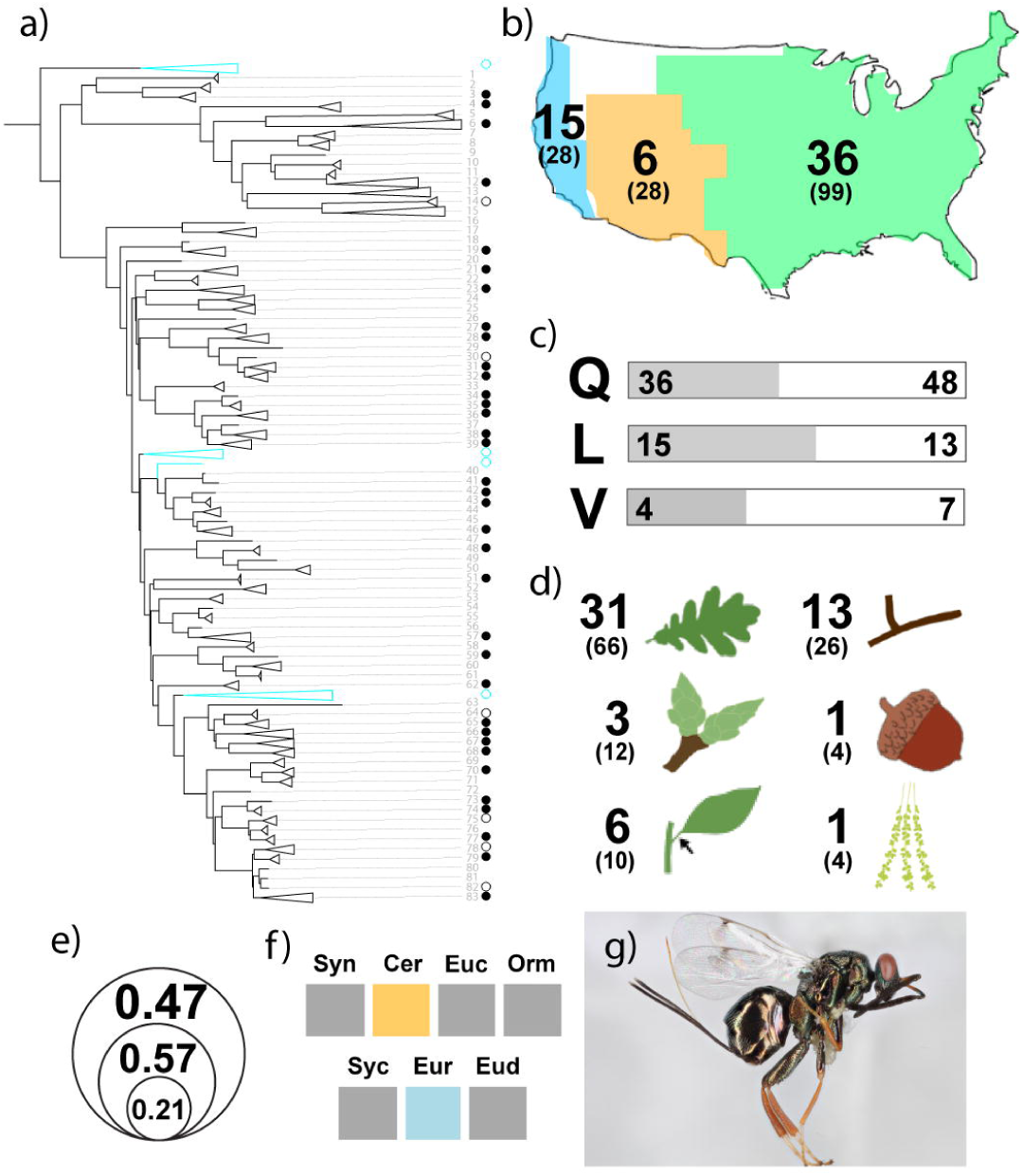
Summary of data for *Torymus* collected from Nearctic galls. For full explanation of figure details, refer to Figure 1 legend. a) Associations of *Torymus* with the Nearctic oak gall wasp phylogeny (Supplementary Figure 1). b) Gall types from which *Torymus* were reared in the Californian (blue), Mexican and Central American (orange), and Eastern North American (green) floristic provinces. c) Associations of *Torymus* with trees in sections Quercus (Q), Lobatae (L), and Virentes (V). d) Association of *Torymus* with gall types on different oak tissues. e) proportion of gall types of three size categories (“small” <0.5 mm; “medium”, “large” > 20 mm) from which *Torymus* were reared. f) Results of probabilistic co-occurrence analysis for *Torymus* against seven other common associates. Yellow = significantly less likely to co-occur; blue = significantly more likely to co-occur; gray = no difference from probabilistic expectations. g) *Torymus* lateral habitus.

#### Biogeography and oak tree section

We reared *Torymus* from galls in all three floristic regions (Figure 5b) and from all three oak sections (Figure 5c). *Torymus* are also known from galls on oaks in section *Protobalanus* in California (Grissell 1976).

#### Tree organ and gall size

*Torymus* wasps were found in association with galls on all six organ types (Figure 5d). They emerged more often from large and medium sized galls than from small (>5 mm) galls (Figure 5e). Though our collections were exclusive to above-ground galls, *Torymus* have also been reared from root galls (Forbes et al. 2016).

#### Co-occurrence with other natural enemies

*Torymus* were reared from the same gall types producing *Eurytoma* wasps significantly more often than predicted (P = 0.0200). Conversely, *Torymus* were significantly less likely to be reared from the same gall types as *Ceroptres* wasps (Figure 5f; P = 0.0003). Previous work has shown that *Torymus* wasps can emerge from the same individual galls as other insects, though these other insects are usually either known or suspected inquilines (e.g., *Synergus*, *Brasema*), or suspected parasites of inquilines (e.g., *Allorhogas* Gahan) (Hall 2001).

#### Additional Notes

The implication from the current *Torymus* taxonomic organization (Grissell 1976) and from molecular phylogeny of family Torymidae (Janšta et al. 2018) is that oak galls have been colonized two or more times, such that gall-associated *Torymus* are para- or polyphyletic. Any future phylogenetic assessment of *Torymus* coevolution with oak galls should therefore endeavor to include non-gall associated taxa.

#### *Sycophila* Walker, 1871 (Hymenoptera: Chalcidoidea: Eurytomidae: Eurytominae)

16,712 individuals (mean = 253.2, range 1 – 9,934) reared from 66 gall types (Supplementary Table 1).

#### Summary of Natural History

*Sycophila* are known as parasitoids of endophytic insects, including gall wasps (Askew et al. 2006; 2013; Balduf 1932; Gómez et al., 2013). Some species are known from just a single host (e.g., *Sycophila marylandica* (Girault) (Balduf 1932), while the Palearctic species *Sycophila buguttata* (Swederus) has 80 recorded hosts (Askew et al. 2013). Recent molecular work has shown considerable cryptic diversity and more limited host ranges among the Nearctic species, (Zhang et al. 2022).

#### Relationship to galler phylogeny

*Sycophila* in our collections were broadly associated with almost all clades in the Nearctic gall wasp phylogeny (Ward et al. 2022) gall wasp phylogeny, including all Palearctic clades (Figure 6a). Two clades from which no *Sycophila* were reared were a mixture of cluster galls on leaves and stems, early spring bud galls, and small monothalamous leaf galls. *Sycophila* appear to be reared more often from large galls (Figure 6e; Hall 2001, Zhang et al. 2022) such that this apparent absence may reflect a general favoring of larger galls, but could also or instead be related to phenology or a bias in survivorship from smaller galls when using our rearing methods.

**Figure 6.**
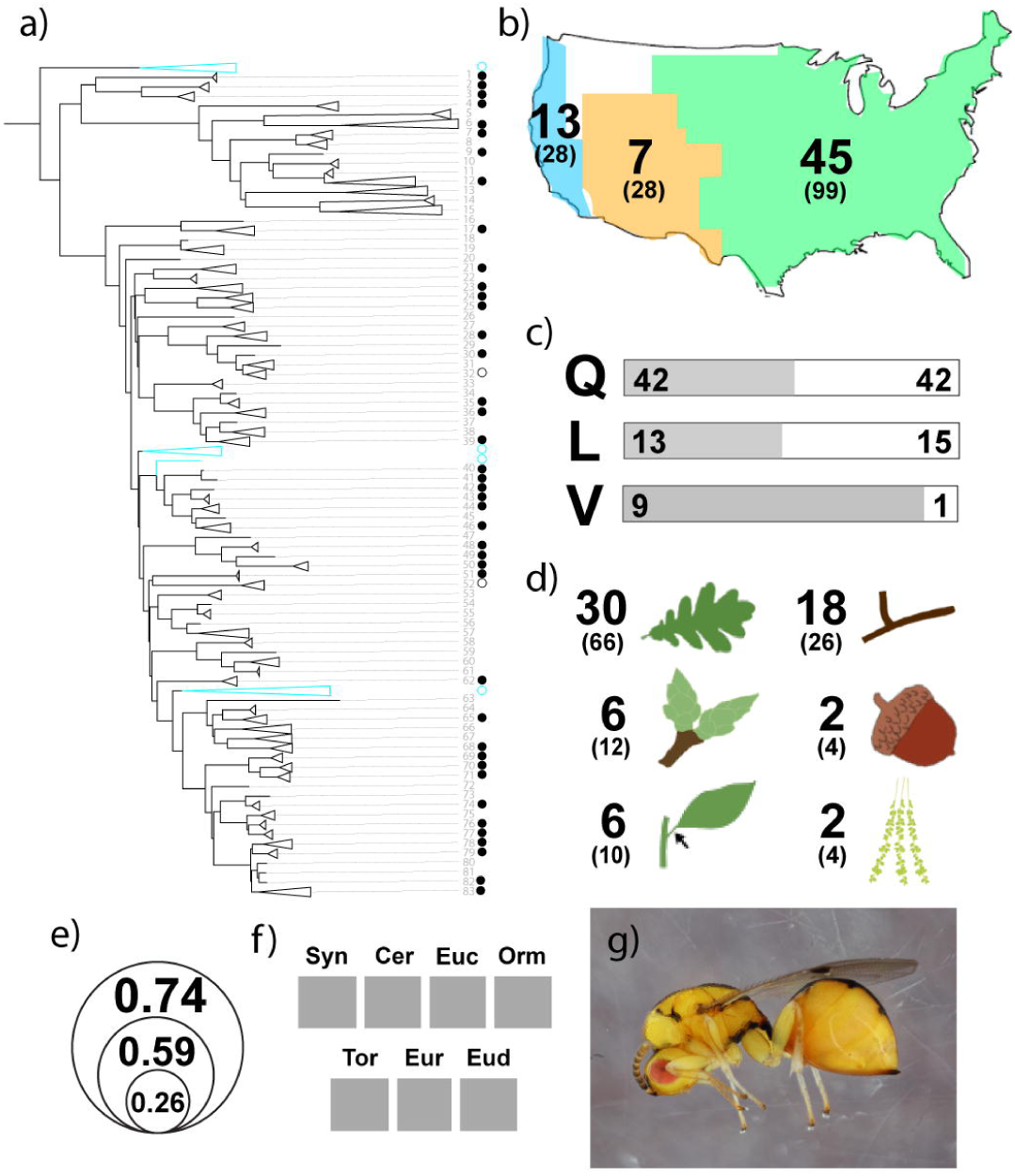
Summary of data for *Sycophila* collected from Nearctic galls. For full explanation of figure details, refer to Figure 1 legend. a) Associations of *Sycophila* with the Nearctic oak gall wasp phylogeny (Supplementary Figure 1). b) Gall types from which *Sycophila* were reared in the Californian (blue), Mexican and Central American (orange), and Eastern North American (green) floristic provinces. c) Associations of *Sycophila* with trees in sections *Quercus* (Q), *Lobatae* (L), and *Virentes* (V). d) Association of *Sycophila* with gall types on different oak tissues. e) proportion of gall types of three size categories (“small” <0.5 mm; “medium”, “large” > 20 mm) from which *Sycophila* were reared. f) Results of probabilistic co-occurrence analysis for *Sycophila* against seven other common associates. Yellow = significantly less likely to co-occur; blue = significantly more likely to co-occur; gray = no difference from probabilistic expectations. g) *Sycophila* lateral habitus.

#### Biogeography and oak tree section

*Sycophila* were reared from galls on oaks across all three regions and in all three oak sections (Figures 6b, c). Nine of 10 galls reared from live oaks (section *Virentes*) were host to *Sycophila*.

#### Tree organ and gall size

We reared *Sycophila* from all surveyed host tree organs (Figure 6d). At least at this genus-level resolution, they were reared from a larger fraction (74%) of large galls than from medium (59%) or small (26%) galls (Figure 6e). PCoA analyses of *Sycophila* from these and other collections also suggest that wasps in the genus generally favor medium and large galls (Zhang et al. 2022).

#### Co-occurrence with other natural enemies

*Sycophila* did not co-occur significantly in a positive or negative direction with any of the other seven genera analyzed (Figure 6f).

#### *Eurytoma* Illiger, 1807 (Hymenoptera: Chalcidoidea: Eurytomidae: Eurytominae)

4,170 individuals (mean = 62.2, range 1 – 1,605) reared from 67 gall types (Supplementary Table 1).

#### Summary of Natural History

*Eurytoma* in the Nearctic comprise a large (> 80 species) group of mostly parasitic wasps associated with a large diversity of insects across several orders (Bugbee 1967). At least 10 Nearctic species have been reared from oak galls, but their direct host is often uncertain, and indeed some Palearctic species are confirmed to attack non-galling inquilines (Redfern and Askew 1998). Some *Eurytoma* are also phytophagous, although no exclusive phytophages are known from oak galls (Bugbee 1967). Though some *Eurytoma* are apparently endoparasites (Redfern and Askew 1998), *Eurytoma* in other Cynipid galls, including oak galls in the Palearctic, are uniformly ectoparasites, with species in some galls feeding on the gall organ once the primary host insect has been exhausted (Gómez et al. 2011).

Reported host ranges of oak gall-associated *Eurytoma* vary from a single species to more than 75 hosts (Bugbee 1967, Gómez et al. 2011; Askew et al. 2013). However, the diversity of the Nearctic fauna has yet to be interrogated genetically, and those in the Palearctic only marginally so. One particularly generalist-appearing species in the Palearctic, *Eurytoma brunniventris* Ratzeburg, has shown some evidence of genetic structure at the COI locus, with specimens reared from five species of oak gall wasps sorting genetically by tree host section (Ács et al. 2002). Until an integrative assessment of species limits can be performed for Nearctic *Eurytoma*, interpretation of their host ranges will likely remain limited.

#### Relationship to galler phylogeny

*Eurytoma* were or have previously been reared from most oak galls represented in the Nearctic gall wasp phylogeny (Ward et al. 2022) (Figure 7a). If they appear sparse anywhere on the phylogeny, it is among the *Neuroterus* part of the tree (Supplementary Figure 1; tips 4- 17), many of which were collected as small flower or leaf galls.

**Figure 7.**
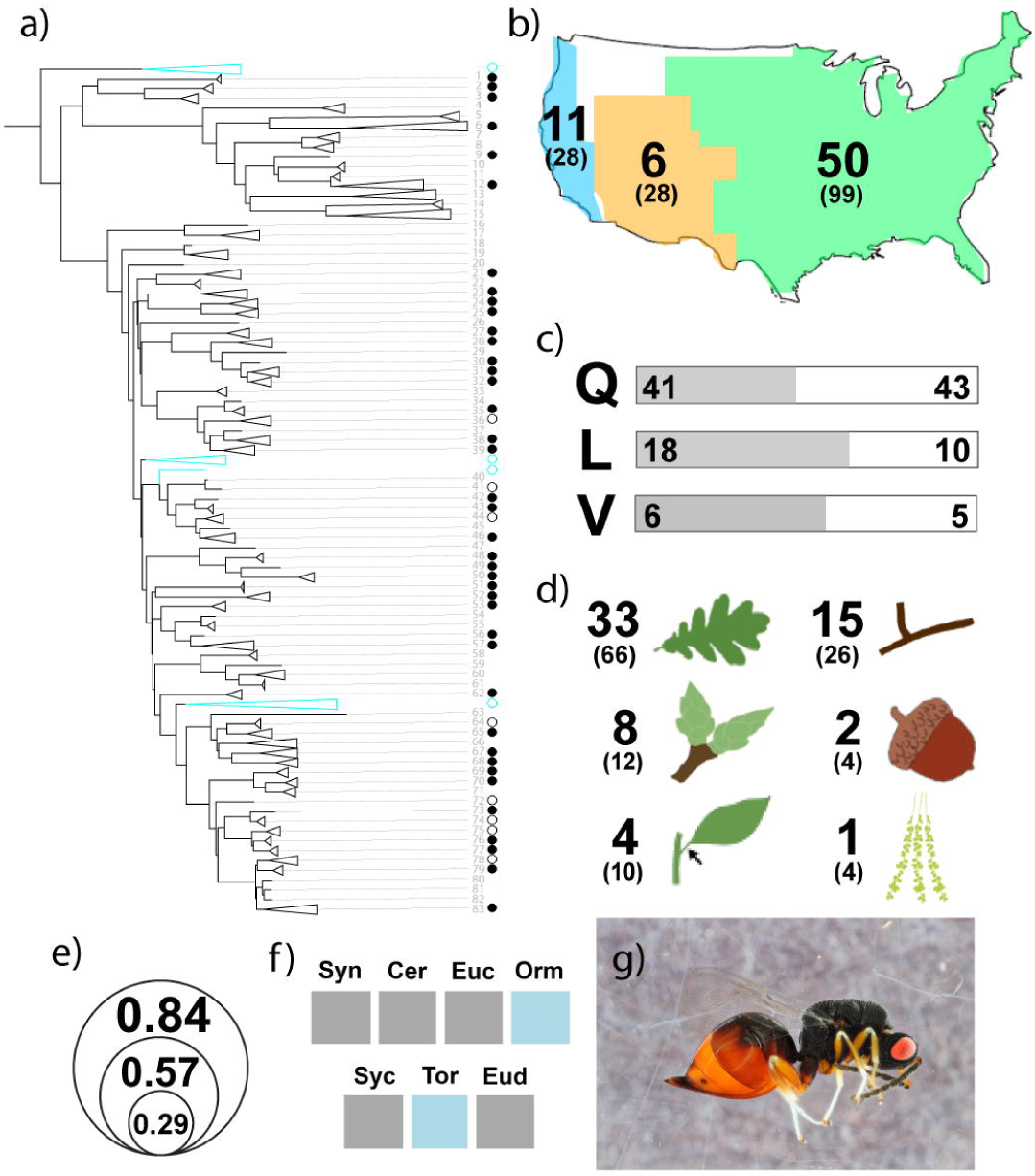
Summary of data for *Eurytoma* collected from Nearctic galls. For full explanation of figure details, refer to Figure 1 legend. a) Associations of *Eurytoma* with the Nearctic oak gall wasp phylogeny (Supplementary Figure 1). b) Gall types from which *Eurytoma* were reared in the Californian (blue), Mexican and Central American (orange), and Eastern North American (green) floristic provinces. c) Associations of *Eurytoma* with trees in sections *Quercus* (Q), *Lobatae* (L), and *Virentes* (V). d) Association of *Eurytoma* with gall types on different oak tissues. e) proportion of gall types of three size categories (“small” <0.5 mm; “medium”, “large” > 20 mm) from which *Eurytoma* were reared. f) Results of probabilistic co-occurrence analysis for *Eurytoma* against seven other common associates. Yellow = significantly less likely to co-occur; blue = significantly more likely to co-occur; gray = no difference from probabilistic expectations. g) *Eurytoma* lateral habitus.

#### Biogeography and oak tree section

*Eurytoma* were reared from galls across all three floristic regions (Figure 7b) and from 48-64% of gall types collected from each oak section (Figure 7c).

#### Tree organ and gall size

*Eurytoma* were reared from galls on all oak organs studied (Figure 7d) though were more often reared from larger galls (84%) than from small galls (29%; Figure 7e).

#### Co-occurrence with other natural enemies

*Eurytoma* were significantly more likely to be present when *Ormyrus* (P = 0.0033) or *Torymus* (P = 0.0200) wasps were also present (Figure 7f). This may indicate overlap in the types of gall morphologies favored by each parasitoid genus.

#### Additional Notes

Like *Ormyrus* and T*orymus*, *Eurytoma* are near-ubiquitous in their association with oak galls: we reared them from 64 (52.5%) of the 122 gall types that had five or more insect specimens emerge (Supplemental table 1). Also like these other genera, *Eurytoma* may be more species-rich than they currently appear, with each species more specialized on particular dimensions of gall environments (Zhang et al. 2014). A thorough integrative analysis of this group will be necessary to address questions about their evolution, ecology, and taxonomy.

#### *Euderus* Haliday 1844 (Hymenoptera: Chalcidoidea: Eulophidae: Entiinae)

816 individuals (mean = 81.6, range 1 – 402) reared from 10 gall types (Supplementary Table 1).

#### Summary of Natural History

*Euderus* is a moderately large genus, with >75 species described worldwide. Where hosts are known, they are usually pupae in concealed habitats (e.g., leaf mines, inside fruits and stems) (Yoshimoto 1971). More rarely, *Euderus* attack insects in galls, with two oak gall associated species – *Euderus crawfordi* Peck and *Euderus set* Egan, Weinersmith & Forbes – known from the Nearctic (Yoshimoto 1971; Egan et al. 2017). *Euderus set* has been specifically studied as a behavioral manipulator of its host gall wasps – wasps with *E. set* infections chew significantly smaller exit holes in their galls than those that are not infected and then do not leave the gall but instead plug the exit hole with their head. *Euderus set* then consumes the body of the host wasp and leaves the gall by chewing a hole through its host’s head (Weinersmith et al. 2017; Ward et al. 2019). Though this behavior has only been studied in detail for *E. set*, evidence of “head plugs” has been found in museum collections of Southwestern U.S. *Bassettia* Ashmead galls (Egan et al. 2017).

#### Relationship to galler phylogeny

We reared or found records of *Euderus* in association with nine gall types in the Nearctic gall wasp phylogeny (Ward et al. 2022) phylogeny. Seven of these (indicated by “s” in Figure 8a) have been previously identified as *E. set* (Ward et al. 2019). *Euderus crawfordi* (“c” in Figure 8a) has been reared from the Palearctic species *Plagiotrochus suberi* Weld, but only from galls in its introduced range in the Nearctic suggesting this is a derived host association. *Euderus crawfordi* is also known from the Nearctic *Kokkocynips coxii* (Bassett), which is not on the Ward et al. (2022) tree, but we have indicated its approximate location near its congener, *Kokkocynips imbricariae* (Ashmead) (“c” with two asterisks). Our collections also produced what appears to be a new species of *Euderus* associated with galls of the sexual generation of *Neuroterus washingtonensis* on the Pacific coast (“n” in Figure Xa). The morphology of these wasps did not match that of *E. set*, nor any species described in Yoshimoto (1971). Two other *Euderus* reared from galls in the southwestern U.S. were not examined morphologically and their hosts were not among those on the Ward et al. (2022) tree.

**Figure 8.**
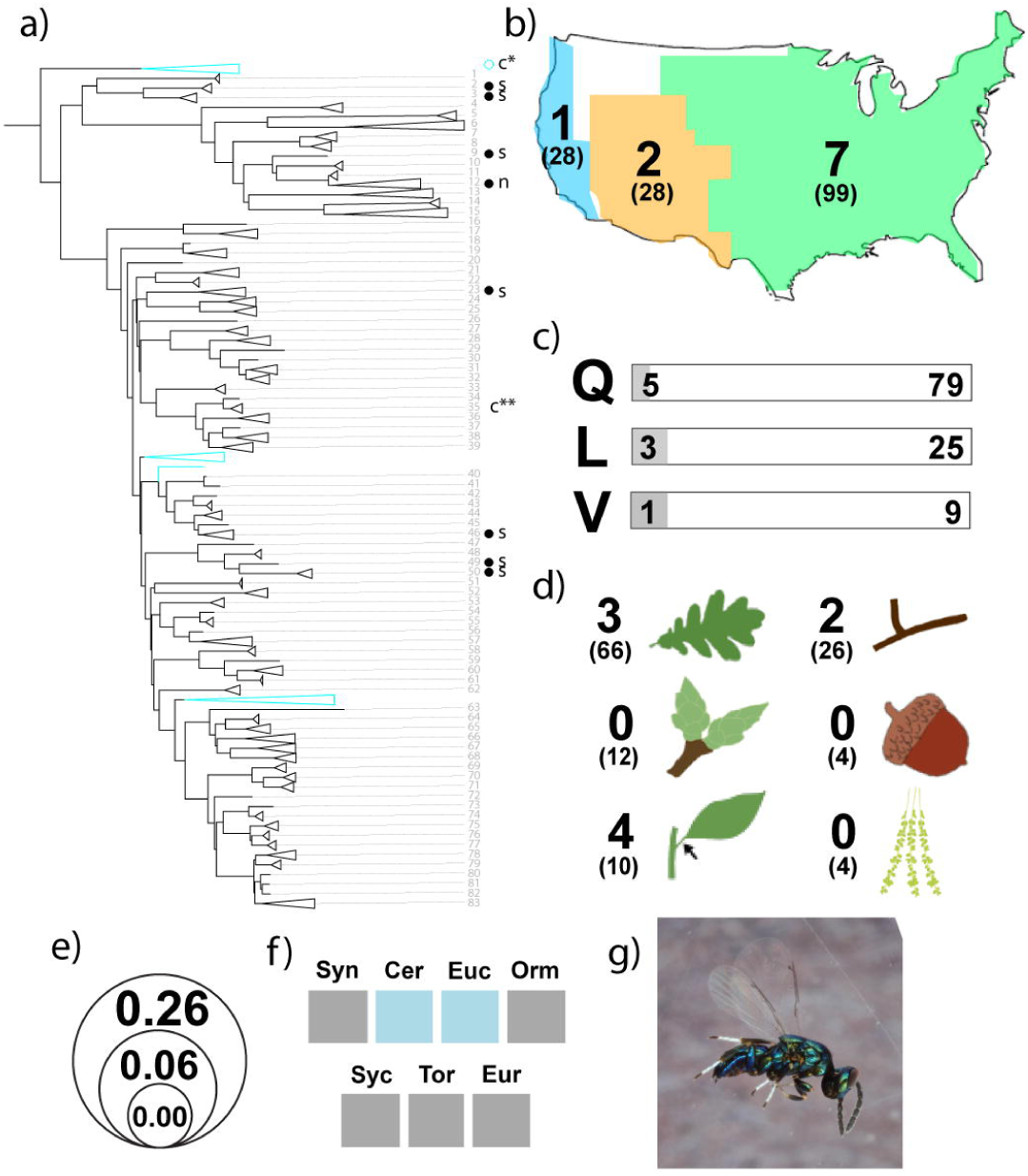
Summary of data for *Euderus* collected from Nearctic galls. For full explanation of figure details, refer to Figure 1 legend. a) Associations of *Euderus* with the Nearctic oak gall wasp phylogeny (Supplementary Figure 1). “s” = *Euderus set*, “c” = *Euderus crawfordi*, “n” = possible new *Euderus* species. * = this association with a Palearctic oak tree is only known from trees introduced to California. ** = the approximate phylogenetic location of this host of *E. crawfordi* (see text). b) Gall types from which *Euderus* were reared in the Californian (blue), Mexican and Central American (orange), and Eastern North American (green) floristic provinces. c) Associations of *Euderus* with trees in sections *Quercus* (Q), *Lobatae* (L), and *Virentes* (V). d) Association of *Euderus* with gall types on different oak tissues. e) proportion of gall types of three size categories (“small” <0.5 mm; “medium”, “large” > 20 mm) from which *Euderus* were reared. f) Results of probabilistic co-occurrence analysis for *Euderus* against seven other common associates. Yellow = significantly less likely to co-occur; blue = significantly more likely to co-occur; gray = no difference from probabilistic expectations. g) *Euderus* lateral habitus.

#### Biogeography and oak tree section

*Euderus set* was reared from seven gall types, all in Eastern North America but across all three oak sections (Figure 8b,c). *Euderus* were reared from two types of gall in the Mexican and Central American floristic region (both on section Quercus oaks), and from one gall type in the Californian floristic province (also section Quercus).

#### Tree organ and gall size

Gall hosts were only on stems, leaves, or petioles (Figure 8d). Most gall hosts were large (>2cm), and no *Euderus* were reared from galls smaller than 5mm (Figure 8e).

#### Co-occurrence with other natural enemies

*Euderus* were found to occur alongside *Ceroptres* (P = 0.015) and *Euceroptres* (P = 0.001) significantly more often than expected (Figure 8f). This co-occurrence is unlikely to be because *Euderus* are using wasps in either of these two genera as their primary hosts - the particular biology of *Euderus set* causes the host to be visible from outside of the gall and the gall inducer has been shown to be the host cases where this has been investigated (Weinersmith et al. 2017; Ward et al. 2019). Other hypotheses for the significant co-occurrence are that wasps in these three genera tend to attack galls with similar features, and/or that they all tend to attack galls in a similar temporal window (Ward et al. 2019).

#### Additional Notes

Because we are primarily working at the level of genus or above and patterns of specialization are more likely to manifest at the species level we have not sought to test hypotheses about insect specialization on gall morphology in this paper. However, since we have *Euderus* rearing records to the level of species, here we can compare features of their associated galls. Previous work has noted that *Euderus set* has only been reared from integral galls that lack external defensive structures, perhaps because *Euderus* appear to attack late-stage pupae and have relatively short ovipositors (Ward et al. 2019). This pattern of host use – medium to large, integral (non-detachable) swellings, often with >1 cells – appears to apply to *E. crawfordi* and to the putative new *Euderus* species reared from *Neuroterus washingtonensis* as well (Figure 8a; Supplemental Table 1).

### Other Common Affiliates - Superfamily and Family level IDs

#### Eulophidae (Hymenoptera: Chalcidoidea: Eulophidae)

We reared 21,232 (mean = 259, range = 1 – 8,655) eulophid wasps from 82 gall types. These counts exclude wasps in subfamily Entiinae (*Euderus*), which were treated separately above. Previously documented Eulophid associates of oak galls include representatives of the Entedoninae (*Chrysochari*s Förster, *Eprhopalotus* Girault, *Horismenus* Walker, *Pediobius* Walker), Eulophinae (*Aulogymnus* Förster, *Cirrospilus* Westwood, *Pnigalio* Schrank, *Sympiesis* Förster), and Tetrastichinae (*Aprostocetus* Westwood, *Baryscapus* Förster, *Galeopsomyia* Girault, *Minotetrastichus* Kostjukov, *Pentastichus* Ashmead, *Quadrastichus* Gilrault, *Tetrastichus* Walker) (Askew et al. 2013, Noyes 2022).

Relatively little is known about the ecology of eulophid gall associates, including their trophic roles, host ranges, and details of their life cycles. Representatives of many eulophid genera and subfamilies are known as obligatory or facultative hyperparasitoids (Schauff et al. 1997), and eulophid larvae in some oak galls can be gregarious (Redfern and Askew 1998). Several different eulophid species have previously been reared from the same gall type (e.g., Askew 1961, Eliason and Potter 2000, Bird et al. 2013), but they may or may not be attacking the same host(s) in these communities. Future integrative taxonomic work that includes host relationships will be invaluable in understanding this important group of gall associates.

#### Eupelmidae (Hymenoptera: Chalcidoidea: Eupelmidae)

We reared 10,620 (mean = 190; range = 1– 3,802) eupelmid wasps from 56 gall types (Supplementary Table 1). The Eupelmidae most commonly associated with oak galls are genera *Brasema* or *Eupelmus* Dalman, and it is likely that all or most of our collections are from one of these genera (though we note that *Anastatus gemmaii* (Ashmead) was originally recorded from *Callirhytis quercusgemmaria* Ashmead (Burks 1967)). Many *Brasema* have been mistakenly classified as *Eupelmus*, and sexual dimorphism in the family makes it challenging to separate males (Gibson 2011), such that we felt it was premature to split the two genera here.

*Brasema* and *Eupelmus* are both ectoparasitic on insect larvae or pupae, and many previous Holarctic oak gall associations have been recorded (Noyes 2022). Associations with Palearctic oak gallers are known to involve direct parasitism of oak gall wasps, as well as parasitism of their *Synergus* inquilines (Redfern and Askew 1998). We have at least two records of *Brasema* emerging from the same monothalamous leaf gall as an adult *Belonocnema kinseyi* Weld oak gall wasp (Hall 2001), strongly suggesting that at least some of our collections represent indirect associations with the galler.

#### Ichneumonoidea (Hymenoptera)

We reared a combined 229 (mean = 11.5, range = 1 – 80) ichneumonid and braconid wasps from 20 gall types. All but one of the ichneumonoid collections were reared from medium-sized galls or larger (>0.5cm) on stems, leaves, or petioles (Supplementary Table 1). Though we did not key all of these wasps to genus, the majority of these collections were wasps in the genus *Allohorgas* (Braconidae: Doryctinae). *Allohorgas* has previously been reared from several North American galls (Eliason and Potter 2000; Forbes et al. 2016; Weinersmith et al. 2020; Joele et al. 2021). Braconid wasps from subfamily Cheloninae were reared from *Disholcaspis quercusmamma* (Walsh & Riley) and *Callirhytis frequens* (Gillette) galls.

The ecology of the oak gall-associated wasps in superfamily Ichneumonoidea is still generally unknown. Recent work on genus *Allorhogas* shows it to be particularly labile with respect to its feeding habits, with some species acting as parasites, others as phytophagous inquilines, still others as seed predators, and some as gall formers themselves (Moreira et al. 2017; Samaca-Saenz et al. 2020, 2022). In oak galls, *Allorhogas* and other ichneumonid wasps may be attacking moth or beetle inquilines: Joseph et al (2011) reported *Bassus nucicola* Musebeck (Braconidae) and an unidentified ichneumonid wasp, both thought to be parasitizing *Cydia latiferreana* Walsingham in galls of *Andricus quercuscalifornicus*. Cheloninae are also primarily known as parasitoids of Lepidoptera (Wharton et al. 1997).

#### Pteromalidae (Hymenoptera: Chalcidoidea: Pteromalidae)

We reared 7,545 (mean = 148; range = 1– 2,385) pteromalid wasps from 51 different gall types (Supplementary Table 1). Our pteromalid collections were reared from galls in all three floristic regions and from all three oak tree sections.

Existing keys, which acknowledge their own difficulty and incompleteness (e.g., Bouč and Heydon 1997), were insufficient for separating many of these wasps to the genus level within this large and polyphyletic family, so they will be addressed elsewhere in the future through integrative taxonomic approaches. The Western Palearctic fauna of oak gall-associated Pteromalidae includes at least nine genera (*Arthrolytus* Thomson*, Cecidostiba* Thomson*, Caenacis* Förster*, Cyrtoptyx* Delucchi*, Elatoides* Nikol’skaya*, Hobbya* Delucchi*, Kaleva* Graham*, Mesopolobus* Westwood, and *Ormocerus* Walker) (Noyes 2022). At least three of these (*Arthrolytus*, *Cecidiostiba*, and *Ormocerus*) are also known from Nearctic oak galls. Additional Nearctic gall-associated pteromalids include *Acaenacis* Girault, *Amphidocius* Dzhanokmen, *Anisopteromalus* Ruschka, *Guolina* Heydon, *Lariophagus* Crawford, and *Pteromalus* Swederus.

Where the ecology of oak gall-associated pteromalids has been studied, they are ectoparasites of a variety of gall inhabitants. Some attack the gall wasp itself, while others parasitize inquilines or other parasitoids (Askew 1961ab). *Mesopolobus* are the best studied of the Palearctic oak gall-associated pteromalids and feed on several different gall inhabitants, and at various life stages, including adults (Askew 1961). As many as five different species from this genus have been reared from the same gall type (Redfern and Askew 1998). Experiments on host searching suggest that short range searches for galls by *Mesopolobus* wasps may rely more on visual cues than on olfaction (Askew 1961), such that host tree and/or gall morphology may be axes of specialization to explore for these and other pteromalids.

### Occasional associates (Hymenoptera)

#### Bethylidae (Hymenoptera: Chrysidoidea: Bethylidae)

We reared 37 (mean = 5, range 1 – 16) bethylid wasps from seven gall types. A previous study found a wasp in bethylid genus *Goniozus* Förster in a gall of *Belonocnema fossoria* Weld (Forbes et al. 2016), and other *Goniozus* have been reared from moth inquilines in oak galls (Fouts 1942). The four gall types with the most bethylid wasps in our collections (*Andricus quercuspetiolicola* and three species of *Belonocnema*) also produced moths, indicating a possible trophic connection.

#### *Bootanomyia* Girault, 1915 (Hymenoptera: Chalchidoidea: Megastigmidae)

We reared 2,385 (mean = 596, range 1 – 2,307) wasps in genus *Bootanomyia* Girault from four gall types, all leaf galls on *Quercus garryana* on the Pacific coast. While several species from this genus are common parasites of oak galls wasps in the Palearctic (Doğanlar 2012, Askew et al. 2013), only one informal record exists of the genus in North America (photos #2020534-6 on BugGuide.org) and these are the first records we can find of *Bootanomyia* associated with oak gall wasps in the Nearctic. Though we did not key all of our collections to species, we keyed one to *Bootanomyia dorsalis* (Fabricius) using Doğ nlar (2012).

Molecular analysis of *B. dorsalis* in the western Palearctic found evidence for host-associated genetic differentiation among wasps reared from oak galls on trees in different oak subgenera (Nicholls et al. 2018). Whether our collections represent an introduced population of *B. dorsalis*, another (or several) host-associated populations, or a combination of these two remains to be seen.

#### Ceraphronoidea (Hymenoptera)

We reared 24 ceraphronoid wasps (mean = 8, range 2 – 11) from three different *Belonocnema* leaf galls. These were extremely uncommon, accounting for <0.15% of all associates reared from either gall type. The biology of most ceraphronoids is generally poorly known (Johnson and Musetti 2004), but some have been reared from galls of cecidomyiid midges (Loiácono and Margaría 2002).

#### Chalcididae (Hymenoptera: Chalcidoidea)

We reared just two chalcidid wasps, one from each of two gall collections: *Belonocnema fossoria* and *Neuroterus washingtonensis*. Most Chalcididae are parasites of Lepidoptera or are sometimes hyperparasites of other Hymenoptera (though usually still in lepidopteran systems; Boucek and Halstead 1997). Our *B. fossoria* collections did have associated Lepidoptera. While we did not officially record lepidopterans from our *N. washingtonensis* collections, we observed larvae and moths emerging from these fleshy galls. Some chalcidids are known from Lepidoptera-induced galls (e.g., Prinsloo 1984), so these records may represent rare oviposition “mistakes.”

#### Crabronidae (Hymenoptera: Apoidea)

We reared three crabronids from galls of *Andricus wheeleri* Beutenmüller collected in Arizona and two more from galls of *Disholcaspis quercusmamma* collected in Minnesota. Some Crabronidae (e..g, Pemphredoninae) create larval cells in hollow spaces associated with plants (Ashmead 1894) and so these may have been occupying a gall emptied of its original inhabitants.

#### Encyrtidae (Hymenoptera: Chalcidoidea)

We reared 22 encyrtid wasps (mean = 3.6, range 1 – 16) from six gall types. Encyrtidae are usually known as endoparasitoids and often are hyperparasitic on other hymenoptera (Noyes et al. 1997). The rarity of these wasps among our collections leads us to believe that these were non-specific attacks on other gall associates.

#### Eumeninae (Hymenoptera: Vespoidea: Vespidae)

We reared nine mason wasps (mean = 3, range 1 – 5) from three gall types. All three galls (*Disholcaspis quercusmamma, Andricus quercuscalifornicus,* and *Disholcaspis quercusglobulus* (Fitch)) were medium to large stem galls that are often retained on oak branches even after gall wasps and other insects have exited. While we did not identify the mason wasps in A. quercuscalifornicus, those in the two *Disholcaspis* galls were Bramble mason wasps (*Ancistrocerus adiabatus* (de Saussure)). *Ancistrocerus adiabatus* create larval cells of mud in existing cavities and provision cells with moth caterpillars. Bramble mason wasps are known to use abandoned homes of other insects, including galleries of cerambycid beetles, nests of other wasps, and empty galls (Gosling 1978; Holm 2021). Joseph et al. (2011) previously recorded an unidentified species of vespid wasp in galls of *Andricus quercuscalifornicus*, and suggested it may be acting as a facultative predator. Also, though they were not part of these collections, yellow jacket wasps in the genus *Vespula* (Hymenoptera: Vespoidea: Vespidae: Vespinae) have been reared from *Callirhytis quercusbatatoides* (Ashmead) on live oaks (SPE, pers. obs.).

#### Formicidae (Hymenoptera: Formicoidea)

We found 70 ants (mean = 14, range 1 – 28) in association with five of our collections: *Tapinoma* Foerster in *Amphibolips confluenta* (Harris) galls, *Camponotus* Mayr (a queen), *Temnothorax* Mayr, and *Tetramorium* Mayr in *Callirhytis quercuscornigera* (Osten-Sacken) galls, *Camponotus* (workers and a queen), *Temnothorax* (workers and a queen), and *Crematogaster* Lund in *Disholcaspis quercusmamma* galls, *Temnothorax* on *Andricus quercusstrobilanus* (Osten-Sacken), *Brachymyrmex patagonicus* Mayr and *Crematogaster ashmeadi* Mayr in *Bassettia pallida* Ashmead (previously reported in Weinersmith et al. 2020).

All ants recovered here are almost certainly colonists of older galls that had already been hollowed out by their original inhabitants. Previous studies have reported other ant species in other galls, e.g., *Camponotus nearcticus* Emery and *Lasius alienus* (Foerster) in galls of *C. quercuscornigera* (Eliason and Potter 2000), and seven different ant genera in galls of *Disholcaspis cinerosa* (Bassett) (Wheeler and Longino 1988).

Besides living inside oak galls, some ant species are known to act as mutualists, feeding on honeydew produced by some gall types while actively defending those galls from parasites and predators (Washburn 1984, Fernandes et al. 1999). We have observed ants tending *Disholcaspis eldoradensis* (Beutenmuller) galls (which produces honeydew) in the Pacific northwest (KMP, pers. obs.) and *Crematogaster* ants both living inside and tending galls of *D. cinerosa* in the southeastern U.S. (SPE, pers. obs.).

#### Platygastridae (Hymenoptera: Platygastroidea)

We reared 913 platygastrids (mean = 91.3, range 1 – 867) from 10 gall types. The vast majority (867) of these were reared from *Neuroterus washingtonensis* galls on the Pacific coast but other sporadic collections came from Eastern galls. Weinersmith et al. (2020) previously reported three platygastrid genera (*Telenomus* Haliday, *Calotelea* Westwood, and *Synopeas* Förster) all from the same gall type. Platygastridae are usually egg parasites and *Synopeas* are known to parasitize cecidomyiid midges (Abram 2012). Midges were reared from six of these 10 gall types (Supplemental Table 1), such that these rearing records might not reflect direct associations with the gall wasp or its parasites.

#### Trichogrammatidae (Hymenoptera: Chalcidoidea)

We reared 64 trichogrammatid wasps (mean = 21.3, range 1 – 62) from three gall types (62 from *Neuroterus vesicula* (Bassett), one each from two other gall types). Trichogrammatidae are parasites of insect eggs (Pinto 1997). A preliminary identification of these wasps keyed them to genus *Poropoea* Förster, known for parasitizing eggs of leaf-rolling weevils. It seems likely that these represent accidental collections of insect eggs and not formal gall associates.

### Occasional associates – other insect orders and non-insect arthropods

#### Coleoptera

We collected 413 adult or larval beetles (mean = 10.9, range 1 – 123) from 38 gall types. We did not identify all beetle collections beyond the level of order, but recognized Curculionidae, Ptinidae, and Staphylinidae among families represented. Many different beetles have previously been found in association with Nearctic galls, including Buprestidae, Cerambycidae, Cleridae, Curculionidae, Latridiidae, and Ptinidae (Eliason and Potter 2000, Joseph et al. 2011, Forbes et al. 2016, Weinersmith et al. 2020). Of these beetle families, all besides Latridiidae are known to bore into galls (or at least into wood) or use galls as shelters (Arnett and Thomas 2000; Sugiura and Yamazaki 2009). When the fates of insects reared from individual galls have been tracked, galls with beetle emergents produce no other insects (e.g., Hall 2001), suggesting that they effectively function as direct or indirect predators of all other gall associates.

#### *Forficula* L., 1758 (Dermaptera: Forficulidae)

Two adult forcifulid earwigs in genus *Forficula* L. (Engel 2003) emerged from galls of *Amphibolips quercusinanis* (Osten-Sacken) one and five days after gall collection. These nocturnal insects may have been using the large, mostly-hollow *A. quercusinanis* galls as a daytime shelter.

#### Cecidomyiinae (Diptera: Cecidomyiidae)

We reared 169 cecidomyiid midges (mean = 7.3, range 1 – 70) from 23 gall types. Though one possibility is that some small number of cecidomyiid galls were collected alongside oak galls, Redfern and Askew (1998) suggest that cecidoymiids may sometimes be part of the successional fauna of cynipid oak galls, using the gall tissue after the original inhabitants have emerged. Previous records of cecidomyiids reared in cynipid oak gall studies include *Lasioptera* Meigen and *Lestodiplosis* Keiffer from *C. quercuscornigera* (Eliason and Potter 2000) and unidentified species reared from *B. kinseyi* and *B. pallida* (Forbes et al. 2016; Weinersmith et al. 2020).

#### *Lonchaea* Fallén 1820 (Diptera: Lonchaeoidea: Lonchaeidae: Lonchaeinae)

We reared 30 Lonchaeinae (lance flies) from galls of *C. quercuscornigera*. These specimens keyed to genus *Lonchaea* Fallén (McAlpine et al. 1987). Lance flies are often associated with fallen trees and burrows of bark beetles (McAlpine et al. 1987, Marshall 2012).

#### *Eidalimus* Kertesz 1914 Pachygastrinae (Diptera: Stratiomyoidea: Stratiomyidae)

We reared 12 pachygastrinids from the same galls as the *Lonchaea* flies above. These specimens keyed to genus *Eidalimus* Kertesz (McAlpine et al. 1981).

#### Aphididae (Hemiptera: Aphidoidea)

We found seven aphids (mean = 2.3, range 1 – 3) in association with three gall types. Two were on leaf galls, the other on a stem gall. We did not key these beyond family. We assume these aphids were feeding on or next to galls and that their collection was coincidental to the presence of the gall.

#### *Orius* Wolff, 1811 (Hemiptera: Cimicoidea: Anthocoridae)

We found four minute pirate bugs (genus *Orius* Wolff) - one adult and three nymphs - across three gall types. *Orius* can be both predaceous and herbivorous. We suspect this was another non-specific gall association.

#### Psyllidae (Hemiptera: Psylloidea)

We found 251 psyllids associated with two galls: 250 from *Callirhytis quercuspunctata* (Bassett) and one from *Andricus incertus* Bassett. All 250 collected from *C. quercuspunctata* were collected from the same city (St. Louis, MO), and 249 of these were from the same collection. Psyllids can be gall formers themselves, but here are likely sap feeders and their collection alongside galls may have been due to generalist feeding or entirely coincidental.

#### Rhyparochromidae (Hemiptera: Lygaeoidea)

We found one rhyparochromid (dirt-colored seed bugs) in a collection of *Disholcaspis mellifica* Weld gall from California. These are seed-feeders, and this collection was likely coincidental.

#### Lepidoptera

We reared 355 moths (mean = 19.7; range 1 – 140) from 18 different gall types. We did not key all moths to family and some of the larger galls collected by the KP/DJ team (Supplemental Table 1; lettered lab codes) were observed to have moths emerge but these insects were not tallied. Moths reared from other galls have often been described as or assumed to be inquilines feeding on gall tissue (Joseph et al. 2011, Forbes et al. 2016). Seventeen of the 18 gall hosts of Lepidoptera in our collections were large or medium galls, and most with thick outer walls which would provide ample food for a larger inquiline species.

Moths in the families Gelechiidae, Pyralidae, Sesiidae, and Tortricidae have been previously reared from Nearctic galls (Eliason and Potter 2000, Joseph et al. 2011, Forbes et al. 2016). Most moths in our collections were small moths (10-20 mm wingspan), but we also reared the clearwing moths *Synanthedon scitula* Harris and *Synanthedon decepiens* Edwards from some *Callirhytis* galls. These larger (up to 30 mm) moths mimic vespid wasps and bees and their larvae feed on wood of living trees.

#### Lacewings (Neuroptera: Chrysopoidea: Chrysopidae)

Eleven lacewings (mean = 1.4, range 1 – 3) were found in association with eight gall types. Of these, ten were nymphs and one was an adult. The adult was associated with a collection of *Disholcaspis quercusmamma* bullet galls and may have been emerged from a pupae inside a hollowed-out gall. Lacewing larvae and adults are generalist predators and may have been coincidental collections. However, on live oaks (section *Virentes*), lacewings have been observed to lay eggs on the asexual fuzzy leaf gall induced by *Andricus quercuslanigera* (SPE, pers. obs). It is not known whether this is coincident to the presence of the gall or if proximity to the gall increases lacewing survivorship in some way.

#### Psocomorpha (Psocodea)

We reared 318 barklice (mean: 18.7, range 1 – 156) from 17 gall types. Barklice in gall systems have previously been regarded as late-stage inquilines or successional associates (Joseph et al. 2011) and have been observed to enter empty galls through the exit holes of other gall inhabitants (Weinersmith et al. 2020).

#### Thysanoptera

We found 62 thrips (mean = 2.2, range 1 – 10) in association with 28 gall types. We did not key these specimens beyond the level of order. Though some thrips can be gall inducers or kleptoparasites of other thrips-induced galls (Crespi and Abbot 1999), our thrips collections may have been occupying old hollow galls (e.g., Redfern and Askew 1998). They also may not have been specifically associating with galls but rather collected accidentally while feeding externally on oak tissue.

#### Acari (Arachnida)

We found 135 mites (mean = 7.1, range 1 – 50) in association with 19 gall types. Eliason and Potter (2000) documented mites in families Oribatidae, Phytoseiidae, and Acaridae on the surface of *C. quercuscornigera* galls or sheltering in crevices. *Histiogaster robustus* Woodring (Acaridae) was also found to be apparently phoretic on *C. quercuscornigera* and its *Synergus* inquilines (Eliason and Potter 2000). In our collections, we also found mites were attached to other gall associates, especially ants and beetles.

#### Araneae (Arachnida)

We found 41 spiders (mean = 5.9, range 1 – 18) in association with collections of seven different gall types. All galls were 5 mm or larger. Spiders were likely collected as transients on external gall material, though conceivably may have been inside empty gall cavities of woody stem galls such as *An. quercuscalifornicus, D. quercusglobulus, and D. quercusmammma.* Eliason and Potter (2000) previously recorded spiders in the families Araneidae, Linyphiidae, Philodromidae, Salticidae, and Theridiidae on galls of *C. quercuscornigera*, and also observed spiders eating gall wasps that had emerged from galls.

#### Pseudoscorpiones (Arachnida)

A single pseudoscorpion was found in association with a *Callirhytis quercusbatatoides* gall collected in Florida. Redfern and Askew (1998) mention pseudoscorpions as successional species in galls.

#### Chilopoda

We found one centipede in a gall of *Amphibolips quercusinanis*. Like the *Forficula* earwigs associated with the same galls, this animal was likely using the gall as a transient shelter.

## Summary and Future Directions

These collections provide an abundance of new information about the inhabitants of Nearctic oak galls. We found that some of the most common genera associated with oak galls are reared from galls across the breadth of the gall wasp phylogeny (e.g., *Synergus*, *Ormyrus*, *Torymus*), while other genera (e.g., *Ceroptres*) appear not to attack certain gall wasp clades. Some less species-rich genera (e.g., *Euceroptres*), as well as some individual species (e.g., *E. set*) also offer a look into apparent patterns of specialization that implicate non-phylogenetic axes of adaptation as being important to host ranges.

With an estimated 700 species of oak gall wasp in the Nearctic (Melika et al. 2021b), most of which likely have two generations with morphologically distinct galls, many more interactions remain to be discovered. Our collections are most complete for the Midwestern United States, leaving other regions only lightly sampled. Our only collections in Canada were from Vancouver Island in British Columbia, and we had no collections from Mexico, where much of the Nearctic oak diversity is concentrated and which is likely the home of considerable undiscovered Nearctic oak gall wasp diversity (Egan et al. 2018, Manos and Hipp 2021). Sampling oak gall wasps and their associates from across their entire geographic, temporal, and ecological ranges in the Nearctic represents a major undertaking, but a necessary one if we hope to fully understand the ecology and evolution of these complex communities.

These and other collections of gall wasp associates should be paired with integrative taxonomic work that includes molecular markers. Recent work in *Ormyrus* (Sheikh et al. 2022), *Synergus* (Ward et al. 2020, Driscoe et al. in prep), and *Sycophila* (Zhang et al. 2022) have shown that current species limits based solely on morphology often lump many species into one group and consequently over-inflate host ranges for those species. Some gall wasp species may themselves harbor cryptic host-associated genetic diversity (Ward et al. 2022). As host ranges are critical for understanding adaptation and evolution in gall-associated insects, accurate delimitation of putative species is of paramount importance.

Another future goal should be the discovery of trophic and food web connections for each of these many gall associates, across many different gall types and gall wasp generations. Rearing studies such as this one are important for establishing many of the players in the system, but understanding their respective roles requires careful ecological study. Gall dissections and study of developing larvae and pupae (e.g., Askew and Redfern 1998) have proved useful in this respect for the Western Palearctic fauna. Molecular methods may also be useful for detection of otherwise obscured endoparasites or parasite eggs that may have been oviposited but were subsequently overcome by the host insect’s immune system (e.g., see methods in Condon et al. 2014). Descriptions of ecological relationships are critical for deciphering patterns of host shifting and coevolution revealed by phylogenetic studies, and for connecting those patterns to underlying mechanisms.

A major goal that synthesizes many of the objectives above should be disentangling the geographic histories and histories of host use evolution for both gall wasps and their insect associates. Gall wasps in the Nearctic have shifted between oak tree hosts - and onto different tree organs - many times (Ward et al. 2022). These patterns, alongside empirical evidence from the model *B. treatae* system (Egan et al. 2012ab, Zhang et al. 2017, 2021ab, Hood et al. 2019), as well as the broader live oak cynipid community (Egan et al. 2013, Zhang et al. 2019), imply that host shifts and geography have both been instrumental in speciation for gall wasps and their associates. Cophylogenetic methods that connect the gall wasp phylogeny to those of each of their common associates will allow for tests of whether and how often gall wasp host shifting might cascade from host to parasite (or host to inquiline, as the case may be).

Phylogenies should also reveal whether different parasite and inquiline genera have parallel histories of host gall associations or whether their paths have been more independent.

In summary, these collection data contribute to the growing set of resources available for studying Nearctic oak gall wasps and their associated insect communities. They provide new biogeographic and ecological context for several gall-associated parasitic and inquilinous genera and underscore the potential of this system for addressing myriad critical research questions.

## Acknowledgements

For their help with collections and rearing, we thank Robin Bagley, Megan Blance, Will Carr, Isaac Carroo, Matt Comerford, Biplabendu Das, Sara Devine, Jose Di Paola, Sarah DeLong-Duhon, Rachel Erikson, Brad Foley, Leo Gastel, Alaine Hippee, Elaine Hu, Rebecca Izen, Omar Khodor, Susan Lee, Sean Liu, Kyle McElroy, John McElroy, Daniel McGarry, Briley Mullin, Kevin Neely, Maurine Neiman, Shannon Pelini, Gaston Del Pino, Pedro Brandão DF Pinto, James Reynolds, Catherine Ruis, Monzer Shakally, Thienthanh Trinh, Eric Tvedte, Joseph Verry, Camilla Vinson, Paul Ward, Paula Ward, Heather Widmayer, Ian Will, and Caleb Wilson. We acknowledge Jeff Clark for his work at Gallformers.org, a resource which has been invaluable to this study. CKD was supported by an NSF REU to AAF (DBI: 1757334). YMZ was supported by Oak Ridge Institute for Science and Education (ORISE) fellowship. KLW was funded by a Faculty Fellowship in Ecology and Evolutionary Biology from Rice University. KMP was funded by grants from the National Geographic Society (NGS-533955R-18) and NSF (DEB 1934387). Mention of trade names or commercial products in this publication is solely for the purpose of providing specific information and does not imply recommendation or endorsement by the USDA. USDA is an equal opportunity provider and employer.

## References Cited

Abe Y, Ide T, Wachi N. 2011. Discovery of a new gall-inducing species in the inquiline tribe Synergini (Hymenoptera: Cynipidae): inconsistent implications from biology and morphology. Ann Entomol Soc Am 104:115–120.

Abram PK, Haye T, Mason PG, Cappuccino N, Boivin G, Kuhlmann U. 2012. Biology of *Synopeas myles*, a parasitoid of the swede midge, Contarinia nasturtii, in Europe. BioControl 57:789–800.

Ács Z, Challis RJ, Bihari P, Blaxter M, Hayward A, Melika G, Csóka G, Pénzes Z, Pujade-Villar J, Nieves-Aldrey J-L, Schönrogge K, Stone GN. 2010. Phylogeny and DNA barcoding of inquiline oak gall wasps (Hymenoptera: Cynipidae) of the Western Palaearctic. Mol Phylogenet Evol 55:210–225.

Arnett Jr RH, Thomas MC (eds.). 2000. American Beetles: Vol 1. Archostemata, Myxophaga, Adephaga, Polyphaga: Staphyliniformia. CRC Press, Boca Raton, FL.

Arnett Jr RH, Thomas MC, Skelley PE, Frank JH (eds.). 2002. American beetles: Vol 2. Polyphaga: Scarabaeoidea through Curculionoidea. CRC Press, Boca Raton, FL.

Ashmead WH. 1887. On the cynipidous galls of Florida, with descriptions of new species and synopses of the described species of North America. Trans Am Entomol Soc 14:125–158.

Ashmead WH. 1894. The habits of the Aculeate Hymenoptera.—I. Psyche 7:19–26.

Ashmead WH. 1903. Classification of the gall-wasps and the parasitic Cynipoids, or the superfamily Cynipoidea III. Psyche 10:140–155.

Askew RR. 1961. On the biology of the inhabitants of oak galls of Cynipidae (Hymenoptera) in Britain. Trans Soc Bri Entomol 14:237–268.

Askew RR. 1965. The biology of the British species of the genus Torymus Dalman associated with galls of Cynipidae on oak, with special reference to alternation of forms. Trans Soc Bri Entomol 16:217–232.

Askew RR, Plantard O, Gomez JF, Nieves MH, Nieves-Aldrey JL. 2006. Catalogue of parasitoids and inquilines in galls of Aylacini, Diplolepidini and Pediaspidini (Hym., Cynipidae) in the West Palaearctic. Zootaxa 1301:1–60.

Askew RR, Melika G, Pujade-Villar J, Schönrogge K, Stone GN, Nieves-Aldrey JL. 2013. Catalogue of parasitoids and inquilines in cynipid oak galls in the West Palaearctic. Zootaxa 3643:1–133.

Bailey R, Schönrogge K, Cook JM, Melika G, Csóka G, Thuróczy C, Stone GN. 2009. Host niches and defensive extended phenotypes structure parasitoid wasp communities. PLoS Biol 7:e1000179.

Balduf WV. 1932. Revision of the chalcid files of the tribe Decatomini (Eurytomidae) in America north of Mexico. Proc US Natl Mus 79:1–95

Bird JP, Melika G, Nicholls JA, Stone GN, Buss EA. 2013. Life history, natural enemies, and management of *Disholcaspis quercusvirens* (Hymenoptera: Cynipidae) on live oak trees. J Econ Entomol 106:1747–1756.

Boucek Z, Halstead J. 1997. Chalcididae. In: Gibson GA, Huber JT, Woolley JB (eds.) Annotated Keys to the Genera of Nearctic Chalcidoidea (Hymenoptera). National Research Council of Canada, Ottawa.

Brookfield JF. 1972. The inhabitants (Hymenoptera: Cynipidae, Chalcidoidea) of the cynipidous galls of *Quercus borealis* in Nova Scotia. Canad Entomol 104:1123–1133.

Bugbee RE. 1967. Revision of chalcid wasps of genus Eurytoma in America north of Mexico. Proc US Natl Mus 118:433–552.

Burks BD. 1967. The North American species of Anastatus Motschulsky (Hymenoptera, Eupelmidae). Trans Am Entomol Soc 93:423–432.

Busbee RW. 2018. Host plant and spatial influences on the natural enemy community structure of a host specific insect herbivore. Texas State University. MS Thesis

Buffington ML, Liljeblad J. 2008. The description of Euceroptrinae, a new subfamily of Figitidae (Hymenoptera), including a revision of *Euceroptres* Ashmead, 1896 and the description of a new species. J Hymenopt Res 17:44–56.

Buffington ML, Forshage M, Liljeblad J, Tang CT, van Noort S. 2020. World Cynipoidea (Hymenoptera): a key to higher-level groups. Insect Syst Div 4:1.

Condon MA, Scheffer SJ, Lewis ML, Wharton R, Adams DC, Forbes AA. 2014. Lethal interactions between parasites and prey increase niche diversity in a tropical community. Science. 343:1240– 1244.

Cooke CL. 2018. Forest Micro-Hymenoptera, including those attacking trees (Cynipidae oak gall wasps) and those potentially defending them (parasitic Pteromalidae). PhD Dissertation, Univ. Maryland.

Cooper WR, Rieske LK. 2011. A native and an introduced parasitoid utilize an exotic gall-maker host. BioControl 56:725–734.

Crespi B, Abbot P. 1999. The behavioral ecology and evolution of kleptoparasitism in Australian gall thrips. Fla Entomol 82:147–164.

Darwin CR. 1875. The variation of animals and plants under domestication. 2nd edition. Volume 2. John Murray, London, UK.

Denk T, Grimm GW. 2010. The oaks of western Eurasia: traditional classifications and evidence from two nuclear markers. Taxon 59:351–366.

Denno RF, McClure MS, Ott JR. 1995. Interspecific interactions in phytophagous insects: Competition reexamined and resurrected. Ann Rev Entomol 40:297–331.

Diehl SR, Bush GL. 1984. An evolutionary and applied perspective of insect biotypes. Annu Rev Entomol 29:471–504

Doğanlar M. 2011. Review of Palearctic and Australian species of *Bootanomyia* Girault 1915 (Hymenoptera: Torymidae: Megastigminae), with description of new species. Turk Entomol Derg 35:123–157.

Drés M, Mallet J. 2002. Host races in plant–feeding insects and their importance in sympatric speciation. Philos Trans R Soc Lond B Biol Sci. 357:471–492.

Driscoe AL, Nice CC, Busbee RW, Hood GR, Egan SP, Ott JR. 2019. Host plant associations and geography interact to shape diversification in a specialist insect herbivore. Mol Ecol 28:4197–4211.

Eady RD. 1959. A revision of the nomenclature in the European Torymidae (Hym. Chalcidoidea) with special reference to the Walker types. — Entomol Mon Mag 94:257–271.

Egan SP, Hood GR, Feder JL, Ott JR. 2012a. Divergent host plant use promotes reproductive isolation among cynipid gall wasp populations. Biol Lett 8:605–608.

Egan SP, Hood GR, Ott JR. 2012b. Testing the role of habitat isolation among ecologically divergent gall wasp populations. Int J Ecol 2012:809897.

Egan SP, Hood GR, DeVela G, Ott JR. 2013. Parallel patterns of morphological and behavioral variation among host-associated populations of two gall wasp species. PLoS ONE 8:e54690.

Egan SP, Hood GR, Martinson E, Ott JR. 2018. Quick Guide: Cynipid gall wasps. Curr Biol 28:PR1370–R1374.

Eliason EA, Potter DA. 2000. Biology of *Callirhytis cornigera* (Hymenoptera: Cynipidae) and the arthropod community inhabiting its galls. Environ Entomol 29:551–559.

Engel MS. 2003. The earwigs of Kansas, with a key to genera north of Mexico (Insecta: Dermaptera). Trans Kans Acad Sci 106:115–123.

Evans D. 1965. The Life History and Immature Stages of *Synergus pacificus* McCracken and Egbert (Hymenoptera: Cynipidae) 1. Can Entomol 97:185–188.

Fagan MM. 1918. The uses of insect galls. Am Nat 52:155–176.

Fernandes GW, Fagundes M, Woodman RL, Price PW. 1999. Ant effects on three-trophic level interactions: plant, galls, and parasitoids. Ecol Entomol 24:411–415.

Fernandes GW, Coelho MS, Santos JC. 2014. Neotropical insect galls: status of knowledge and perspectives. *In*: Fernandes GW, Santos JC (eds.) Neotropical insect galls. Springer, Dordrecht.

Forbes AA, Powell TH, Stelinski LL, Smith JJ, Feder JL. 2009. Sequential sympatric speciation across trophic levels. Science 323:776–779.

Forbes AA, Hood GR, Hall MC, Lund J, Izen R, Egan SP, Ott JR. 2016. Parasitoids, hyperparasitoids, and inquilines associated with the sexual and asexual generations of the gall former, *Belonocnema treatae* (Hymenoptera: Cynipidae). Ann Entomol Soc Am 109:49–63.

Forbes AA, Devine SN, Hippee AC, Tvedte ES, Ward AKJ, Widmayer HA, Wilson CJ. 2017. Revisiting the particular role of host shifts in initiating insect speciation. Evol 71:1126–1137.

Fouts RM. 1942. Description of a new species of *Goniozus* from Oregon (Hymenoptera: Bethylidae). Proc Entomol Soc Wash 44:168–169.

Gallformers Contributors. 2021. Gallformers.org. https://www.gallformers.org. Accessed 25 Apr. 2022.

Gibson GA. 2011. The species of *Eupelmus* (Eupelmus) dalman and *Eupelmus* (Episolindelia) Girault (Hymenoptera: Eupelmidae) in North America north of Mexico. Zootaxa 2951:1–97.

Gibson GA, Huber JT, Woolley JB, Woolley JB. (eds). 1997. Annotated keys to the genera of Nearctic Chalcidoidea (Hymenoptera). NRC Research Press, Ottawa.

Gillette CP. 1896. A monograph of the genus Synergus. Trans Am Entomol Soc 23:85–100.

Gómez JF, Nieves-Aldrey JL, Nieves MH, Stone GN. 2011. Comparative morphology and biology of terminal instar larvae of some *Eurytoma* (Hymenoptera, Eurytomidae) species parasitoids of gall wasps (Hymenoptera, Cynipidae) in western Europe. Zoosystema 33:287–323.

Gómez JF, Nieves-Aldrey JL, Stone GN. 2013. On the morphology of the terminal-instar larvae of some European species of Sycophila (Hymenoptera: Eurytomidae) parasitoids of gall wasps (Hymenoptera: Cynipidae). J Nat Hist 47:2937–2960.

Gosling DC. 2017. Observations on the biology of the oak twig pruner, *Elaphidionoides parallelus*, (Coleoptera: Cerambycidae) in Michigan. Gt Lakes Entomol 11:1.

Goulet H, Huber JT. 1993. Hymenoptera of the world: and identification guide to families. Canada Communication Group - Publishing, Ottawa.

Griffith DM, Veech JA, Marsh CJ. 2016. Cooccur: probabilistic species co-occurrence analysis in R. J Stat Softw 69:1–17.

Grissell EE. 1976. A revision of western Nearctic species of Torymus Dalman (Hymenoptera, Torymidae). Univ of California Press, Berkeley.

Grissell EE. 1995. Toryminae (Hymenoptera: Chalcidoidea: Torymidae) a redefinition, generic classification, and annotated world catalog of species. Int Mem Entomol 2:1–470.

Hall MC. 2001. Community structure of parasitoids attacking leaf galls of Belonocnema treatae on Quercus fusiformis. Texas State University, MS thesis.

Hanson P. 1992. The Nearctic species of *Ormyrus* Westwood (Hymenoptera: Chalcidoidea: Ormyridae). J Nat Hist 26:1333–1365.

Hearn J, Blaxter M, Schönrogge K, Nieves-Aldrey J-L, Pujade-Villar J, Huguet E, Drezen J-M, Shorthouse JD, Stone GN. 2019. Genomic dissection of an extended phenotype: Oak galling by a cynipid gall wasp. PLoS Genetics 15:e1008398.

Hipp AL, Manos PS, González-Rodríguez A, Hahn M, Kaproth M, McVay JD, Avalos SV, Cavender-Bares J. 2018. Sympatric parallel diversification of major oak clades in the Americas and the origins of Mexican species diversity. New Phytol 217:439–452.

Holm H. 2021. Wasps: their biology, diversity, and role as beneficial insects and pollinators of native plants. Pollination Press LLC, Minnetonka, MN.

Hood GR, Forbes AA, Powell TH, Egan SP, Hamerlinck G, Smith JJ, Feder JL. 2015. Sequential divergence and the multiplicative origin of community diversity. Proc Nat Acad Sci USA 112:E5980–E5989.

Hood GR, Zhang L, Hu EG, Ott JR, Egan SP. 2019. Cascading reproductive isolation: Plant phenology drives temporal isolation among populations of a hostL pecific herbivore. Evol 73:554–568.

Ide T, Kusumi J, Miura K, Abe Y. 2018. Gall inducers arose from inquilines: phylogenetic position of a gall-inducing species and its relatives in the inquiline tribe Synergini (Hymenoptera: Cynipidae). Ann Entomol Soc Am 111:6–12.

Janšta P, Cruaud A, Delvare G, Genson G, Heraty J, Kř žková B, Rasplus J-Y. 2018. Torymidae (Hymenoptera, Chalcidoidea) revised: molecular phylogeny, circumscription and reclassification of the family with discussion of its biogeography and evolution of life-history traits. Cladistics 34:627–651.

Joele FR, Zaldívar-Riverón A, Penteado-Dias AM. 2021. Six new species of *Allorhogas* (Hymenoptera, Braconidae, Doryctinae) from south and southeast Brazil with host-plant record. J Hymenopt Res 82:199–220.

Johnson NF, Musetti L. 2004. Catalog of systematic literature of the superfamily Ceraphronoidea (Hymenoptera). Contr Am Entomol Inst 33:1–149.

Joseph MB, Gentles M, Pearse IS. 2011. The parasitoid community of *Andricus quercuscalifornicus* and its association with gall size, phenology, and location. Biodivers Conserv 20:203–216.

Kaartinen R, Stone GN, Hearn J, Lohse K, Roslin T. 2010. Revealing secret liaisons: DNA barcoding changes our understanding of food webs. Ecol Entomol 35:623–638.

Kinsey AC. 1923. The gall wasp genus *Neuroterus* (Hymenoptera). Indiana Univ Stud 10:1–150.

Kinsey AC. 1930. The gall wasp genus Cynips. A study in the origin of species. Indiana Univ Stud 16:1–577.

Krombein KV, Hurd PD, Smith DR, Burks BD. 1979. Catalog of Hymenoptera in America north of Mexico (Vol. 1). Smithsonian Institution Press, Washington D.C.

Lobato-Vila I, Pujade-Villar J. 2019. Revision of world Ceroptresini (Hymenoptera: Cynipidae) with the description of a new genus and five new species. Zootaxa 4685:1–67.

Lobato-Vila I, Cibrián-Tovar D, Barrera-Ruíz UM, Equihua-Martínez A, Estrada-Venegas EG, Buffington ML, Pujade-Villar J. 2019. Review of the *Synergus* Hartig species (Hymenoptera: Cynipidae: Synergini) associated with tuberous and other tumor-like galls on oaks from the New World with the description of three new species from Mexico. Proc Entomol Soc Wash 121:193–255.

Loiácono MS, Margaría CB. 2002. Ceraphronoidea, Platygastroidea and Proctotrupoidea from Brazil (Hymenoptera). Neotrop Entomol 31:551–560.

Manos PS, Hipp AL. 2021. An Updated Infrageneric Classification of the North American Oaks (*Quercus* Subgenus *Quercus*): Review of the Contribution of Phylogenomic Data to Biogeography and Species Diversity. Forests 12:786.

Marshall SA. 2012. Flies: the natural history & diversity of Diptera. Firefly Books, Buffalo, NY.

Martinson EO, Werren JH, Egan SP. 2021. Tissue-specific gene expression shows a cynipid wasp repurposes oak host gene networks to create a complex and novel parasiteL pecific organ. Mol Ecol doi: 10.1111/mec.16159.

McAlpine JF, Peterson BV, Shewell GE, Teskey HJ, Vockeroth JR, Wood DM. 1981. Manual of Nearctic Diptera. Volume 1. Agriculture Canada, Ottawa, Canada.

McAlpine JF, Peterson BV, Shewell GE, Teskey HJ, Vockeroth JR, Wood DM. 1987. Manual of Nearctic Diptera. Volume 2. Agriculture Canada, Ottawa, Canada.

McCracken MI, Egbert DB. 1922. California Gall-making Cynipidae: With Descriptions of New Species (Vol. 3). Stanford University Press, Redwood City.

Melika G, Thuróczy C, Melika G, and Abrahamson WG. 2002. Review of the world genera of oak cynipid wasps (Hymenoptera: Cynipidae, Cynipini). *In*: Melika G, Thuróczy C (eds). Parasitic wasps: evolution, systematics, biodiversity and biological control. Agroinform, Budapest.

Melika G, Pujade-Villar J, Nicholls JA, Cuesta-Porta V, Cooke-McEwen C, Stone GN. 2021a. Three new Nearctic genera of oak cynipid gall wasps (Hymenoptera: Cynipidae: Cynipini): *Burnettweldia* Pujade-Villar, Melika & Nicholls, *Nichollsiella* Melika, Pujade-Villar & Stone, *Disholandricus* Melika, Pujade-Villar & Nicholls; and re-establishment of the genus *Paracraspis* Weld. Zootaxa 4993:1–81.

Melika G, Nicholls JA, Abrahamson WG, Buss EA, Stone GN. 2021b. New species of Nearctic oak gall wasps (Hymenoptera: Cynipidae, Cynipini). Zootaxa 5084:1–131.

Miller RS. 1967. Pattern and process in competition. Adv Ecol Res 4:1–74.

Moreira GR, Eltz RP, Pase RB, Silva GT, Bordignon SA, Mey W, Goncalves GL. 2017. *Cecidonius pampeanus*, gen. et sp. n.: an overlooked and rare, new gall-inducing micromoth associated with *Schinus* in southern Brazil (Lepidoptera, Cecidosidae). ZooKeys 695:37.

Nicholls JA, Schönrogge K, Preuss S, Stone GN. 2018. Partitioning of herbivore hosts across time and food plants promotes diversification in the *Megastigmus dorsalis* oak gall parasitoid complex. Ecol Evol 8:1300–1315.

Noyes JS. 2022. Universal Chalcidoidea Database. http://www.nhm.ac.uk/chalcidoids. Accessed 25 Apr 2022.

Noyes JS, Woolley JB, Zolnerowich G. 1997. Encyrtidae. *In*: Gibson GAP, Huber JT, Wooley JB (eds). Annotated keys to the genera of Nearctic Chalcidoidea (Hymenoptera). NRC Research Press, Ottawa, Canada.

Nylander JA. 2004. Bayesian phylogenetics and the evolution of gall wasps. Doctoral dissertation. Acta Universitatis Upsaliensis.

Pénzes Z, Tang C-T, Bihari P, Bozsó M, Schwéger S, Melika G. 2012. Oak associated inquilines (Hymenoptera, cynipidae, Synergini). TISCIA Monogr Ser. 11:1–66.

Pénzes Z, Tang C-T, Stone GN, Nicholls JA, Schweger S, Bozso M, Melika G. 2018. Current status of the oak gall wasp (Hymenoptera: Cynipidae: Cynipini) fauna of the Eastern Palaearctic and Oriental Regions. Zootaxa 4433:245–289

Pinto JD. 1997. Trichogrammatidae. *In*: Gibson GAP, Huber JT, Wooley JB (eds). Annotated keys to the genera of Nearctic Chalcidoidea (Hymenoptera). NRC Research Press, Ottawa, Canada.

Price PW, Fernandes GW, Waring GL. 1987. Adaptive nature of insect galls. Environ Entomol 16:15–24.

Prinsloo GL. 1984. A new species of *Hockeria* Walker (Hymenoptera: Chalcididae), parasitic in two gall-inciting Lepidoptera from the Namib desert, South West Africa. Afr Entomol 47:239–244.

Prior KM, Hellmann JJ. 2013. Does enemy loss cause release? A biogeographical comparison of parasitoid effects on an introduced insect. Ecol 94:1015–1024.

Pujade-Villar J, Bellido D, Segu G, Melika G. 2001. Current state of knowledge of heterogony in Cynipidae (Hymenoptera, Cynipoidea). Sess Conjunta Entomol 11:87–107.

Redfern M, Askew RR. 1998. Plant galls. Richmond Publishing. Richmond, England.

Ronquist F, Liljeblad J. 2001. Evolution of the gall wasp-host plant association. Evol 55:2503–2522.

Russo RA. 2021. Plant Galls of the Western United States. Princeton University Press. Princeton, NJ, USA.

Samacá-Sáenz E, Egan SP, Zaldívar-Riverón A. 2020. Species Diversity in the Braconid Wasp Genus *Allorhogas* (Doryctinae) Associated With Cynipid Galls on Live Oaks (*Quercus*: Fagaceae) Using Natural History, Phylogenetics, and Morphology. Insect Syst Div 4:3.

Samacá-Sáenz E, Santos BF, Martínez JJ, Egan SP, Shaw SR, Hanson PE, Zaldívar-Riverón A. 2022. Ultraconserved elements-based systematics reveals evolutionary patterns of host-plant family shifts and phytophagy within the predominantly parasitoid braconid wasp subfamily Doryctinae. Mol Phylogenet Evol 166:107319.

Shauff ME, LaSalle J, Coote LC. 1997. Eulophidae. *In*: Gibson GAP, Huber JT, Wooley JB (eds). Annotated keys to the genera of Nearctic Chalcidoidea (Hymenoptera). NRC Research Press, Ottawa, Canada.

Sheikh SI, Ward AKG, Zhang YM, Davis CK, Zhang L, Egan SP, Forbes AA. 2022. *Ormyrus labotus* Walker (Hymenoptera: Ormyridae): another generalist that should not be a generalist is not a generalist. Insect Syst Div 8:1–14.

Smith MA, Eveleigh ES, McCann KS, Merilo MT, McCarthy PC, Van Rooyen KI. 2011. Barcoding a quantified food web: crypsis, concepts, ecology and hypotheses. PLoS One 6:e14424.

Stone GN, Schönrogge K, Crawley MJ, Fraser S. 1995. Geographic and between-generation variation in the parasitoid communities associated with an invading gallwasp, *Andricus quercuscalicis* (Hymenoptera: Cynipidae). Oecologia 104:207–217.

Stone GN, Schönrogge K, Atkinson RJ, Bellido D, Pujade-Villar J. 2002. The population biology of oak gall wasps (Hymenoptera: Cynipidae). Ann Rev Entomol 47:633–668.

Stone GN, Schönrogge K. 2003. The adaptive significance of insect gall morphology. Trends Ecol Evol 18:512–522.

Sugiura S, Yamazaki K. 2009. Gall-attacking behavior in phytophagous insects, with emphasis on Coleoptera and Lepidoptera. Terr Arthropod Rev 2:41–61.

Tooker JF, Helms AM. 2014. Phytohormone dynamics associated with gall insects, and their potential role in the evolution of the gall-inducing habit. J Chem Ecol 40:742–753.

Veech JA. 2013. A probabilistic model for analysing species co-occurrence. Glob Ecol Biogeogr 22:252–260.

Ward AK, Khodor OS, Egan SP, Weinersmith KL, Forbes AA. 2019. A keeper of many crypts: a behaviour-manipulating parasite attacks a taxonomically diverse array of oak gall wasp species. Biol Lett 15:20190428.

Ward, A. K., S. I. Sheikh, and A. A. Forbes. 2020. Diversity, Host Ranges, and Potential Drivers of Speciation Among the Inquiline Enemies of Oak Gall Wasps (Hymenoptera: Cynipidae). Insect Syst. Divers. 4: 3.

Ward AKG, Bagley RK, Egan SP, Hood GR, Ott JR, Prior KM, Sheikh SI, Weinersmith KL, Zhang L, Zhang YM, Forbes AA. 2022. Speciation in Nearctic oak gall wasps is frequently correlated with changes in host plant, host organ, or both. bioRxiv doi: https://doi.org/10.1101/2022.02.11.480154

Washburn JO. 1984. Mutualism between a cynipid gall wasp and ants. Ecol 65:654–656.

Weinersmith KL, Liu SM, Forbes AA, Egan SP. 2017. Tales from the crypt: a parasitoid manipulates the behaviour of its parasite host. Proc R Soc B: Biol Sci 284:20162365.

Weinersmith KL, Forbes AA, Ward AKG, Brandão-Dias PFP, Zhang YM, Egan SP. 2020. Arthropod community associated with the asexual generation of *Bassettia pallida* Ashmead (Hymenoptera: Cynipidae). Ann Entomol Soc Am 113:373–388.

Weis AE, Abrahamson WG. 1986. Evolution of host-plant manipulation by gall makers: ecological and genetic factors in the Solidago-Eurosta system. Am Nat 127:681–695.

Weld LH. 1921. American gallflies of the family Cynipidae producing subterranean galls on oak. Proc US Nat Mus 59:187–246.

Weld LH. 1957. Cynipid galls of the Pacific Slope. Ann Arbor MI Priv. Print.

Weld LH. 1959. Cynipid galls of the eastern United States. Ann Arbor MI Priv. Print.

Weld LH. 1960. Cynipid galls of the southwest. Ann Arbor MI Priv. Print.

Weld LH. 1952. Cynipoidea (Hym.) 1905-1950, being a supplement to the Dalla Torre and Kieffer monograph. Ann Arbor MI Priv. Print.

Wharton RA, Marsh PM, Sharkey MJ. 1997. Manual of the New World genera of the family Braconidae (Hymenoptera) International Society of Hymenopterists, Special Publication. Washington, D.C.

Wheeler J, Longino JT 1988. Arthropods in live oak galls in Texas. Entomol News 99:25–29.

Yee WL. 2008. Host plant use by apple maggot, western cherry fruit fly, and other *Rhagoletis* species (Diptera: Tephritidae) in central Washington state. Pan-Pac Entomol 84:163–178.

Yee WL, Goughnour RB. 2008. Host plant use by and new host records of apple maggot, western cherry fruit fly, and other *Rhagoletis* species (Diptera: Tephritidae) in western Washington state. Pan-Pac Entomol. 84:179–193.

Yoshimoto CM. 1971. Revision of the genus *Euderus* of America north of Mexico (Hymenoptera: Eulophidae). Can Entomol 103:541–578.

Zhang L, Driscoe A, Izen R, Toussaint C, Ott JR, Egan SP. 2017. Immigrant inviability promotes reproductive isolation among host-associated populations of the gall wasp *Belonocnema treatae*. Entomologia Experimentalis et Applicata 162:379–388.

Zhang L, Hood GR, Ott JR, Egan SP. 2019. Temporal isolation between sympatric host plants cascades across multiple trophic levels of host-associated insects. Biol Lett 15:20190572.

Zhang L, Hood GR, Roush AM, Shzu S, Comerford MS, Ott Jr, Egan SP. 2021a. Asymmetric, but opposing reductions in immigrant viability and fecundity promote reproductive isolation among host-associated populations of an insect herbivore. Evol 75:476–489.

Zhang L, Hood GR, Carroo I, Ott JR, Egan SP. 2021b. Context-dependent reproductive isolation: Host plant variability drives fitness of hybrid herbivores. Am Nat 197:732–739.

Zhang YM, Gates MW, Shorthouse JD. 2014. Testing species limits of Eurytomidae (Hymenoptera) associated with galls induced by *Diplolepis* (Hymenoptera: Cynipidae) in Canada using an integrative approach. Can Entomol 146:321–334.

Zhang YM, Sheikh SI, Ward AKG, Forbes AA, Prior K, Stone GN, Gates M, Egan S, Zhang LN, Davis CK, Weinersmith K, Melika G, Lucky A. 2022. Delimiting the cryptic diversity and host preferences of Sycophila parasitoid wasps associated with oak galls using phylogenomic data. bioRxiv. doi: https://doi.org/10.1101/2022.01.21.477213.

