## Supplemental Figure 1 for "The arthropod associates of 155 North American cynipid oak galls"

Supplemental Figure 1. Phylogeny of several Nearctic gall wasps (adapted from Ward et al. 2022) used in Figures 1-8. Numbers are used to assist in referencing tips and clades in manuscript text. “P” indicates a gall wasp is Palearctic in origin.

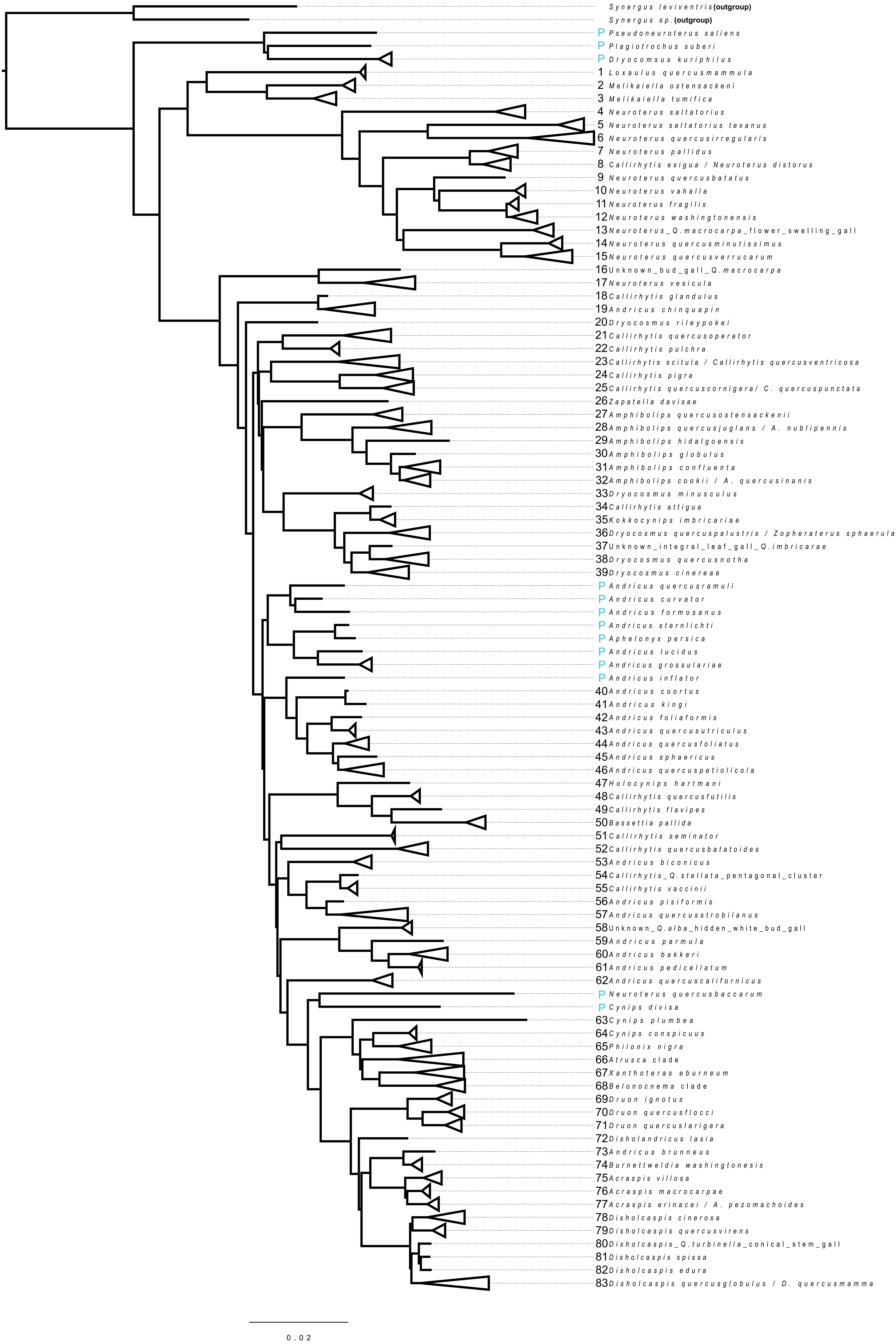
